# Krüppel Regulates Cell Cycle Exit and Limits Adult Neurogenesis of Mushroom Body Neural Progenitors in *Drosophila*

**DOI:** 10.1101/2025.03.24.645006

**Authors:** Dongni Shao Chen, Jin Man, Xian Shu, Haoer Shi, Xue Xia, Yusanjiang Abula, Yuu Kimata

## Abstract

In many organisms, including *Drosophila* and humans, neural progenitors exit the cell cycle and are eliminated by the end of development, thereby restricting adult neurogenesis to specific brain regions. Here, we identify the evolutionarily conserved transcription factor Krüppel (Kr) as a lineage-specific regulator of cell cycle exit and elimination of mushroom body neuroblasts (MBNBs), which generate the learning and memory centre of the *Drosophila* brain, a structure functionally analogous to the mammalian hippocampus. Neuroblast-specific *Kr* RNAi and the *Irregular facet* (*Kr^If-1^*) mutation prolong MBNB lifespan, enabling continued neurogenesis in the adult brain. Although Kr is expressed only at low levels in postembryonic MBNBs, its pupal stage-specific depletion or misexpression is sufficient to cause MBNB retention, revealing a previously unrecognised postembryonic function distinct from its established role in embryonic neurogenesis. Mechanistically, persistent MBNBs maintain expression of the early temporal factor IGF2 mRNA-binding protein (Imp) and fail to fully induce the late temporal factors Syncrip (Syp) and Eip93F (E93). Co-depletion of Imp suppresses MBNB retention caused by Kr depletion, demonstrating that Imp is a key downstream effector of Kr. In parallel, Krüppel homolog 1 (Kr-h1), another Kr family transcription factor and a well-established mediator of hormone-responsive transcription, functionally antagonises Kr by suppressing E93 expression: Kr-h1 knockdown partially rescues the *Kr* depletion phenotype, whereas Kr-h1 overexpression drives tumour-like neuroblast overgrowth. Together, our findings establish Kr as an MBNB-specific coordinator that integrates intrinsic temporal programmes with extrinsic signalling pathways to coordinate neural stem cell termination and neuronal fate transitions, with potential parallels in other organisms.

## Introduction

The adult brain is predominantly composed of postmitotic neurons, which lack the capacity for proliferation. This limitation makes brain tissue particularly vulnerable to damage or degeneration, as lost neurons are largely irreplaceable. Consequently, neuronal loss significantly contributes to the decline of brain function, underlying various chronic pathologies and degenerative diseases associated with aging, such as Alzheimer’s and Parkinson’s diseases (1, 2). Although a small population of proliferative progenitor cells persists in the adult brain, these cells are sparse and primarily quiescent, significantly limiting the brain’s overall regenerative capacity (1, 3).

Neural progenitors are essential for brain development, generating the vast majority of the brain’s cellular constituents. These cells typically undergo asymmetric cell divisions to self-renew and produce differentiated progeny that mature into neurons or glial cells, or remain as partially differentiated intermediate progenitors (4, 5). However, the majority of neural progenitors are eliminated during the late stages of development, leaving behind only a small pool in specialised niches like the subgranular zone in the hippocampus. The remaining progenitors are usually slow-proliferating and dormant but capable of reactivation in response to physiological or pathological stimuli (1, 3). While the elimination of neural progenitors is vital for preserving existing neural circuits, keeping a small subset of progenitors is necessary for repairing brain injury or refining and remodelling neural pathways that support key function during adulthood, such as learning and memory (3, 6, 7). The mechanisms enabling this region-specific control of neurogenic capacity, however, remain poorly understood.

*Drosophila melanogaster* serves as an exemplary model organism for studying the development and regulation of neural progenitors in the central nervous system. In *Drosophila*, neuroblasts (NBs) are responsible for generating virtually all cells in the brain. NBs predominantly undergo asymmetric cell division to generate a daughter NB and a ganglion mother cell, which further divides once to become neurons or glia (5, 8). The neurogenic capacity of NBs, as well as their cell cycle control and survival, are tightly regulated in a lineage/region-specific manner during development (9, 10). At the end of embryogenesis, while many NBs in the abdominal and thoracic segments undergo apoptosis, the majority in the central brain enter a phase of quiescence, reactivated only by larval hatching and subsequent food intake (8, 11, 12). An exception to this regulation is the mushroom body neuroblast (MBNB) lineage, which is responsible for forming the mushroom body, a critical centre for learning and olfactory memory in insects, functionally analogous to the hippocampus in mammals (13, 14). MBNBs continue dividing during the embryo-to-larva transition (9, 15). Furthermore, while other NBs cease dividing and undergo apoptosis or differentiation well before the end of pupal development, MBNBs persist significantly longer until late pupal stages just prior to termination (9, 16, 17). Notably, in some insects like the house cricket *Acheta domesticus* and the moth *Agrotis ipsilon*, MBNBs not only surpass their usual developmental limits but also maintain active neurogenesis throughout the adult phase (18, 19).

Intricate interactions between cell-intrinsic transcription programmes and extracellular cues control the neurogenic capacity of NBs. Temporal transcription factors (tTFs), exemplified by the well-characterised sequence of Hunchback (Hb), Krüppel (Kr), Pdm (Nubbin/Pou domain), Castor (Cas), and Grainyhead (Grh) in embryonic thoracic and many Type I NB lineages, are expressed in NBs in a sequential cascade during development, conferring temporal identity essential for generating the diversity of neural lineages in the central nervous system (20-22). Mammalian neural progenitors may also undergo temporal patterning, as seen in the sequential progression of retinal progenitors and radial glia competence during cortical development (21, 23).

In addition to providing temporal identity, tTFs also regulate NB proliferation and cell cycle progression during development. In embryonic NBs, early tTFs, such as Hb and Kr, promote NB proliferation, while later factors, including Cas and Svp, contribute to limiting NB divisions (11, 24). The transition from early to late tTFs is therefore an important determinant of the duration of neurogenesis and the timing of NB termination (24, 25). In particular, Cas and Svp also influence NB proliferation and lifespan during postembryonic development, and their manipulation can prolong NB proliferation into adulthood. In postembryonic NBs, temporal progression involving Cas and Svp influences NB proliferation and lifespan. For example, Cas overexpression or svp mutation can prolong NB proliferation, allowing NBs to persist and continue generating neural progeny into adulthood (11). In addition to tTFs, two RNA-binding proteins, IGF-II mRNA Binding Protein (Imp) and Syncrip (Syp), also regulate NB cell cycle exit and neuronal fate transitions during postembryonic development. These proteins are expressed in opposing gradients, where Imp promotes NB proliferation and early neuronal fates, whereas Syp promotes NB termination and the transition to late neuronal fates (26, 27). Importantly, while the Imp-Syp gradient is a common regulatory unit among NBs, the regulation of this gradient varies between NB lineages. In Type I NBs, Svp promotes the Imp-to-Syp transition through ecdysone signalling via induction of the Ecdysone Receptor (EcR) (28, 29). In contrast, in MBNBs, where this transition is substantially delayed, activin/Baboon signalling, activated by Myoglianin from surrounding glial cells, regulates this transition (30). Additionally, ecdysone-induced transcription factor Eip93F (E93), a key effector of metamorphic ecdysone signalling, specifically promotes MBNB termination by enhancing autophagy during late pupal stages (31). However, how these intrinsic temporal programmes and extrinsic developmental signals are integrated to regulate lineage-specific termination of MBNBs remains poorly understood.

The Krüppel-like family of transcription factors (KLFs) are conserved across metazoans and play a pivotal role in development, physiology, and stem cell regulation (32, 33). *Drosophila* Kr, a founding member of this family, was first identified as a gap gene, playing a crucial role in establishing the anterior-posterior axis during embryonic development (34). While its critical role as a tTF in embryonic neurogenesis has been recognised (24), the involvement of Kr in postembryonic neurogenesis and beyond has been unexplored. Another KLF member, Krüppel homologue 1 (Kr-h1), is recognised as a target gene of hormonal signalling (35-37). Kr-h1 has been implicated in hormone-dependent neuronal remodelling and is a well-established antagonist of E93 (38, 39), suggesting that it may influence developmental programmes controlling neural progenitor termination. However, the NB lineage-specific roles of Kr and Kr-h1 and their functional interactions remain uncharacterised.

In this study, we uncover a novel function of Kr in regulating the survival and proliferative capacity of MBNBs in *Drosophila* adult brains. We show that NB-specific depletion of Kr, as well as the *Irregular facet* (*Kr^If-1^*) mutation, a classic mutation that causes morphological defects in adult eyes via Kr misexpression (40, 41), extends MBNB lifespan and maintains neurogenesis in the adult brain. We demonstrate that, distinct from its role as a tTF in embryonic NBs, Kr promotes the transition from the Imp-high to the Syp-high state while supporting E93 expression to ensure timely cell cycle exit and elimination of MBNBs during late pupal development. In parallel, we identify an opposing role for Kr-h1, which antagonises E93 and delays MBNB termination. Together, our findings suggest that Kr integrates intrinsic temporal programmes with extrinsic signalling pathways to coordinate neural stem cell termination and neuronal fate transitions, highlighting critical roles of KLF transcription factors in regulating the longevity and neurogenic potential of specific neural progenitor pools.

## Results

### Postmitotic state in the *Drosophila* adult central brain

Early studies concluded that no adult neurogenesis occurs in *Drosophila* adult brains (9, 16, 17). However, a subsequent study employing a highly sensitive lineage tracing technique demonstrated that slow neurogenesis continues beyond adulthood in optic lobes, which can be accelerated by acute injury (42). Additionally, quiescent progenitor cells have been identified in the central brain, capable of inducing neurogenesis or gliogenesis when stimulated by physical damage or excessive neuron death (43, 44). However, under physiological conditions, proliferating cells have not been reported in the adult central brain.

To confirm this, we examined the presence of proliferating cells by immunostaining for the mitotic marker phosphorylated histone H3 Serine 10 (pH3) and by performing EdU incorporation assays to detect DNA synthesis. In control adult brains after five days of EdU feeding, we observed neither pH3-positive cells nor any EdU incorporation (Fig. 1A, Fig. 2A, B). We also examined cell cycle states in brain tissue using Fly-FUCCI, a fluorescent reporter system that visualises cell cycle phases via two differentially fluorescently labelled protein fragments that undergo cell cycle-dependent proteolysis (Fig. 1A) (45). Virtually all cells in the central brain showed only EGFP-E2F1 without mRFP-NLS-CycB signals, indicating that they were in G1 or G0 phase (Fig. 1A), confirming the absence of actively cycling progenitors. Occasionally, one or two cells displaying both EGFP and mRFP signals were observed near the antennal lobes (Fig. 1A). These may correspond to G2 cells previously reported in the adult brain during the so-called “critical period,” which spans five days from eclosion (7, 46).

**Figure 1.**
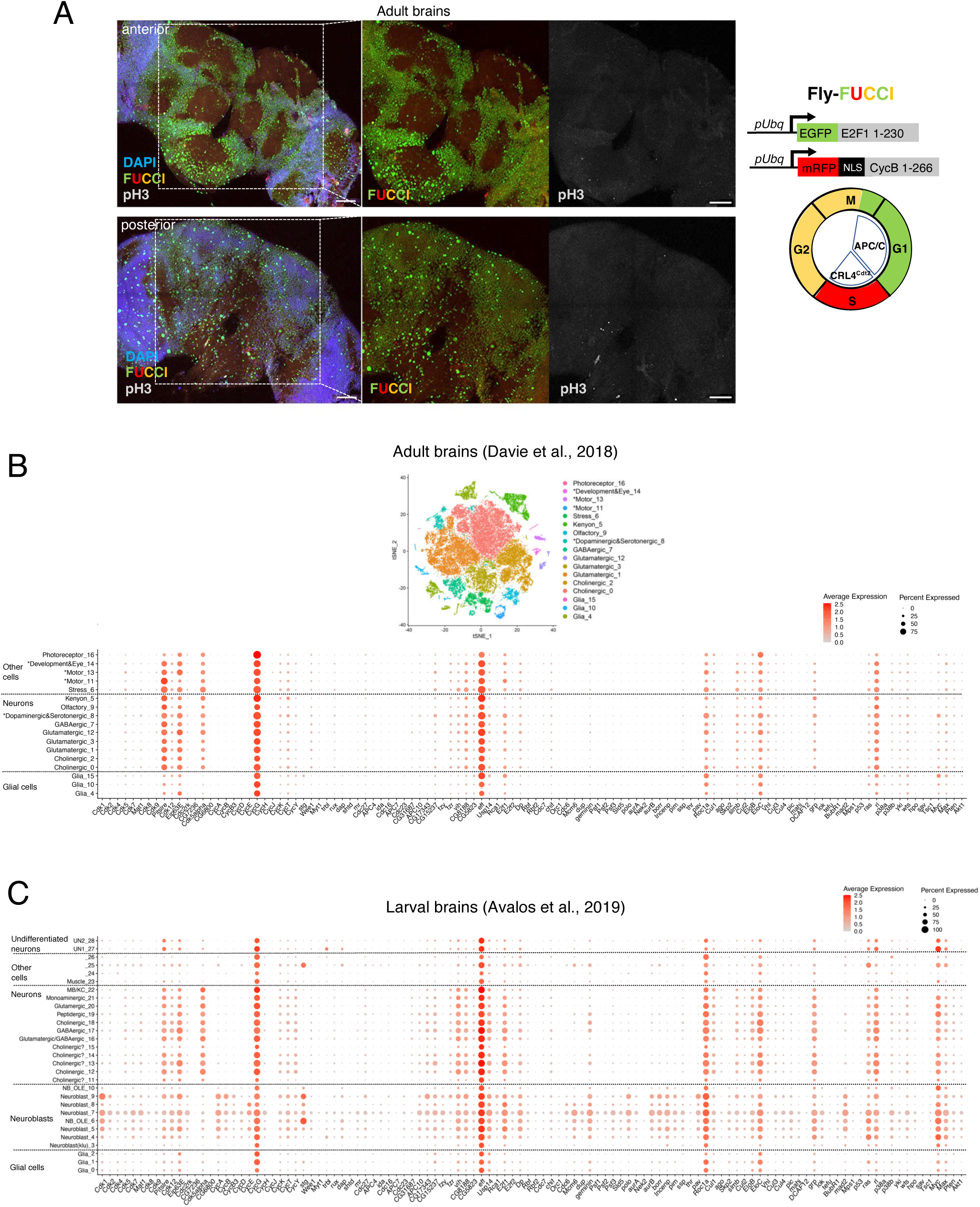
The adult *Drosophila* central brains are post-mitotic. **(A)** Left: representative confocal images of adult control (*w*^1118^) central brains ubiquitously expressing Fly-FUCCI cell-cycle reporters. EGFP::E2F1(1–230) (green) marks cells in G1/G0 phase, whereas mRFP::NLS-CycB (1–266) (red) marks cells in S/G2/M phases. Brains were counterstained with DAPI (blue) and the mitotic marker phosphorylated histone H3 Ser10 (pH3, grey). Upper panels show anterior views and lower panels show posterior views. Nearly all cells in the adult central brain were GFP-positive and lacked detectable RFP signals, indicating that they reside in G1/G0 phase. No pH3-positive mitotic cells were detected. A small number of cells near the antennal lobes exhibited weak RFP signals, suggesting rare G2-phase cells. Scale bars: 50 μm. Right: schematic representation of the Fly-FUCCI system used to visualise cell-cycle states based on cell-cycle-dependent degradation of E2F1 and CycB fragments. **(B)** Top: tSNE plot showing cell clusters identified from adult *Drosophila* brain scRNA-seq data (Davie et al., 2018). Cells are classified into 17 clusters, with different colours indicating neuronal, glial, and neuroblast-like populations. Bottom: dot plot showing expression of 112 selected cell-cycle regulator (CCR) genes across the 17 clusters. Colour intensity indicates average expression level, and dot size indicates the percentage of cells within each cluster expressing the gene. CCR genes are arranged according to broad functional categories. **(C)** Dot plot showing expression of the same selected CCR genes across 29 cell clusters identified from larval *Drosophila* brain scRNA-seq data (Avalos et al., 2019). Colour intensity and dot size are as described in (B). Most positive CCRs are enriched in larval neuroblast clusters but are minimally expressed in adult neuronal and glial clusters, consistent with the postmitotic state of the adult brain.

**Figure 2.**
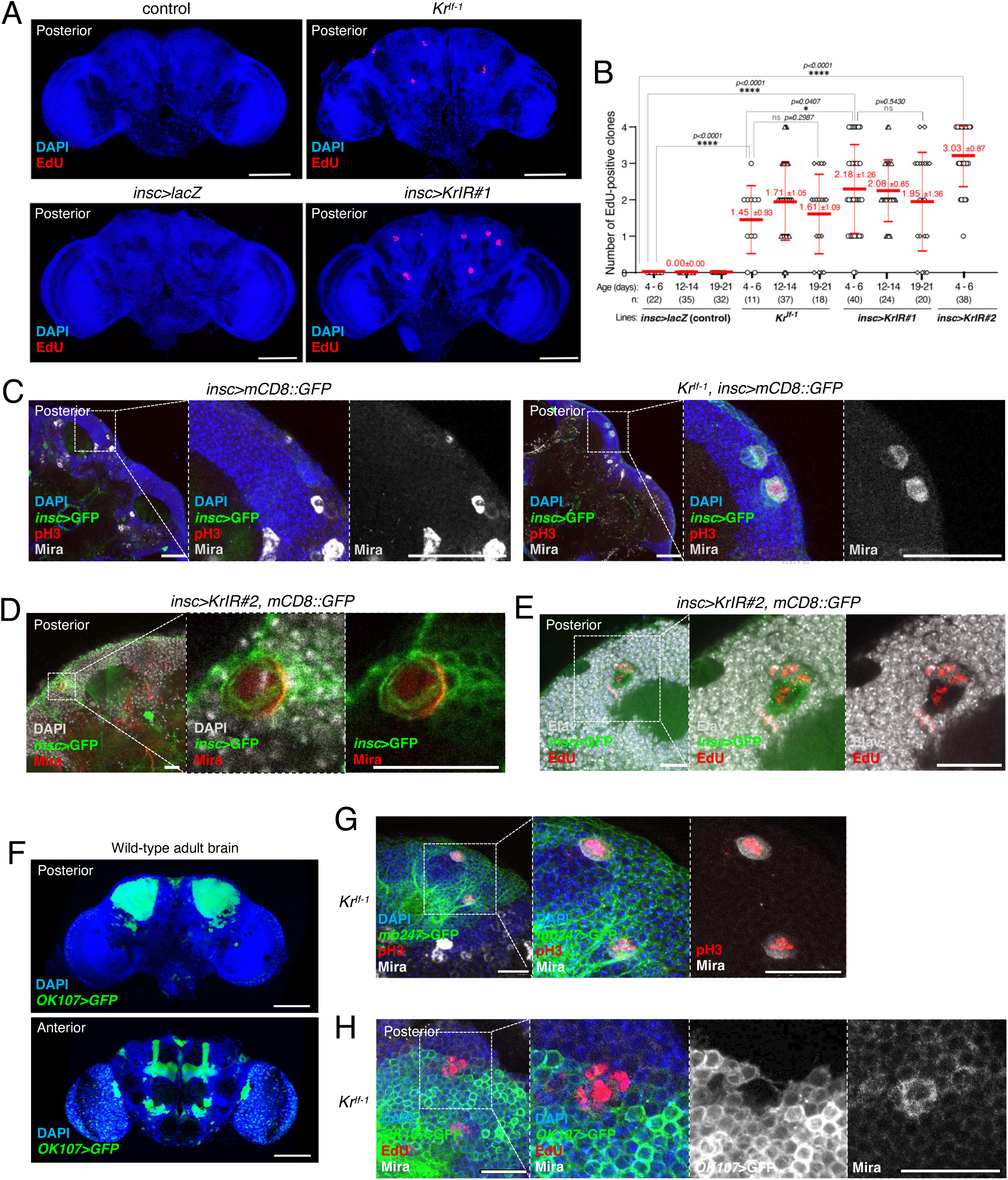
Neuroblast-specific Kr depletion and the *Kr^If-1^* mutation cause mushroom body neuroblast retention and prolonged neurogenesis in adult brains. **(A)** Representative confocal images of adult central brains after EdU incorporation assays. Brains from *Kr* wild-type controls (*Kr*^+^/*Kr*^+^), heterozygous *Kr^If-1^* mutants (*Kr^If-1^*/*Kr*^+^), UAS control flies (*insc>lacZ*) and Kr-depleted flies (*insc>KrIR#1*) were stained with DAPI (blue) and EdU (red). No EdU-positive clones were detected in control brains, whereas *insc>KrIR#1* and *Kr^If-1^* mutant brains contained multiple EdU-positive clones, predominantly in the dorsoposterior region. Scale bars: 100 µm. **(B)** Quantification of EdU-positive clones in adult brains aged 4–6, 12–14 and 19–21 days after eclosion. Scatter dot plots show the number of EdU-positive clones per brain hemisphere. Thick and thin red bars indicate means and SDs, respectively, with values annotated above. n indicates the number of brain hemispheres analysed. Statistical significance was determined using the Mann-Whitney U test. \**p* ≤ 0.05, \*\**p* ≤ 0.01, \*\*\**p* ≤ 0.001, \*\*\*\**p* < 0.0001; ns, not significant. **(C)** Representative confocal images of *Kr* wild-type control and *Kr^If-1^* mutant adult brains expressing mCD8::GFP (green) under the NB-specific *insc*-Gal4 driver. Brains were stained with DAPI (blue), pH3 (red) and Mira (grey). Mitotic and interphase NBs, identified by Mira and GFP expression, were detected in the dorsoposterior region of *Kr^If-1^* mutant brains but not in controls. Scale bars: 50 µm. **(D)** Representative confocal images of *insc>KrIR#2* adult brains expressing mCD8::GFP (green), stained with DAPI (grey) and Mira (red). *insc>GFP*- and Mira-positive mitotic NBs exhibited crescent-like cortical Mira localisation, characteristic of dividing NBs, whereas surrounding smaller cells lacked NB marker expression. Scale bars: 20 μm. **(E)** Representative confocal images of *insc>KrIR#2* adult brains expressing mCD8::GFP (green), stained for EdU (red) and the pan-neuronal marker Elav (grey). EdU-positive cells were observed among Elav-positive neuronal progeny, indicating continued neurogenesis in adult brains following Kr depletion. Scale bars: 20 µm. **(F)** Representative images of wild-type adult brain expressing *mCD8::GFP* (green) under *OK107*-Gal4 driver to visualise MB structures. Brains were stained with DAPI (blue). Posterior and anterior views are shown. The MB cell body region is located dorsoposteriorly, whereas MB lobes are visible in the anterior view. Scale bars: 100 µm. **(G)** Representative confocal images of *Kr^If-1^* mutant adult brains expressing mCD8::GFP under *mb247*-Gal4 (green), stained with DAPI (blue), pH3 (red) and Mira (grey). Mitotic cells in the MB cell body region showed cortical Mira localisation, while surrounding smaller cells lacked NB markers, consistent with dividing MBNBs undergoing asymmetric division. Scale bars: 20 µm. **(H)** Representative confocal images of *Kr^If-1^* adult brains expressing mCD8::GFP under *OK107*-Gal4 after EdU labelling. EdU incorporation was detected in Mira-positive MBNBs, which were weakly marked by *OK107*>GFP. Scale bars: 20 µm.

To gain molecular insight into the postmitotic state of the adult brain, we analysed the expression patterns of cell cycle regulator (CCR) genes using recent single-cell RNA sequencing (scRNAseq) data from *Drosophila* adult and larval brains (47, 48). Cells were organised into 17, 84, or 29 distinct clusters, respectively, based on their gene expression profiles, and the transcription levels of 112 selected CCRs were analysed in each cluster (Fig. 1B, C, Fig. S1). Most CCRs that positively regulate cell cycle progression, including CDKs, cyclins, Polo, Aurora kinases and DNA replication factors, were minimally expressed across all neuronal and glial clusters in both adult and larval brains, while being highly expressed in NB clusters in larval brains (Fig. 1B, C, Fig. S1). A notable exception is E2F1-Dp, a transcriptional activator complex driving the G1/S transition, as well as Myc and Max, a heterodimeric transcription factor that promotes both cell proliferation and growth through transcriptional regulation, which maintained relatively high expression levels in various neuronal and glial clusters (Fig. 1B, C, Fig. S1). In mammals, E2F1-Dp has been implicated in regulating neuronal migration and apoptosis (49-51), while Myc has been associated with neuronal differentiation and activity (52, 53). Thus, E2F1-Dp and Myc-Max complexes may also serve neuron-specific functions in *Drosophila*.

In contrast, negative CCRs, which function to delay or block cell cycle progression, remained relatively high across both NB and neuronal clusters (Fig. 1B, C and Fig. S1). These include components of the anaphase promoting complex/cyclosome (APC/C), such as *fizzy-related* (*fzr,* the *Drosophila* FZR1 orthologue), *vihar* (*vih,* the UBE2C orthologue), CG15237 (the APC15 orthologue), and CG8188 (the Ube2S orthologue), as well as the DNA licensing inhibitor *geminin* and the checkpoint kinase *grape* (*grp*, the CHK1 orthologue).

Besides these canonical CCRs, non-canonical CDKs that function in transcriptional regulation in addition to cell cycle control (54), such as Eip63E/CDK10, Pitslre/Cdk11, and Cdk12, along with their partners CycG, CycK, CycT, and CycY, were relatively highly expressed across both neural and NB clusters (Fig. 1B, C, Fig. S1). Similarly, several CCRs involved in ubiquitin-dependent protein degradation, such as *effete* (*eff,* the *Drosophila* Ube2d2 orthologue), as well as components of E3 ligase complexes, APC/C, SCF, and CRL2, and the deubiquitinase Usp14, exhibited modest to high expression across neural and NB clusters (Fig. 1B, C, Fig. S1), highlighting the importance of transcriptional control and the ubiquitin-proteasome pathway in neuronal regulation (55). A particularly notable CCR was the Cdk5-Cdk5α complex (the CDK5-p35 orthologue), which exhibited significantly higher expression in neural clusters than in NB clusters, supporting its established role in postmitotic neurons (56) (Fig. 1B, C). In summary, our transcriptome analysis suggests that most canonical CCRs are likely to contribute to maintaining the postmitotic state of the adult brain, whereas a subset may perform neuron- and/or glia-specific functions through non-canonical, cell cycle-independent mechanisms.

The above results suggest that repression of positive CCRs, rather than upregulation of negative CCRs, is critical for maintaining the postmitotic state of adult brain tissue. We therefore tested whether forced expression of positive CCRs could drive postmitotic neurons into the cell cycle by overexpressing key positive CCRs in postmitotic mushroom body and dopaminergic neurons using *mb247*-Gal4 and *TH*-Gal4. Co-expression of E2F1-Dp and Cdk2-CycE, but not Cdk4-CycD or CycE and String/CDC25, indeed allowed mitotic entry in both neuron types when expression was induced in young adult flies (less than two days post-eclosion) (Fig. S2A). However, when it was induced 10 days after eclosion, no mitotic neurons were observed (n ≥ 10). Notably, these mitotic neurons frequently exhibited Dcp-1 signals, indicative of apoptosis (Fig. S2B), suggesting that mature neurons become increasingly refractory to cell cycle re-entry with age and that forced entry leads to cell death, possibly through DNA damage accumulation or mitotic defects. We also assessed the physiological impact of this aberrant mitotic entry, conducting a survival analysis, and found that adult flies with E2F1-Dp and Cdk2-CycE induction in dopaminergic neurons exhibited significantly shorter lifespans compared to controls, reducing the median lifespan from 22 days to 14 days (Fig. S2C). Together, these findings indicate that repression of positive CCRs is required to maintain the postmitotic state of adult brain tissue, which is likely to be crucial for normal neuronal function and organismal fitness.

### Neuroblast-specific depletion and mutation of Krüppel cause persistence of proliferative mushroom body neuroblasts in the adult brain

Having confirmed the postmitotic state of the adult *Drosophila* central brain, our next objective was to identify regulators that establish and maintain this state, focusing on NBs. We therefore focused on Krüppel (Kr), a conserved zinc finger transcription factor with established roles in developmental neurogenesis, for functional analysis.

Strikingly, when Kr was depleted specifically in NBs using the *insc*-Gal4 driver and *Kr* RNAi lines, proliferative, EdU-positive clones were observed in the dorsoposterior region of the central brains in adult flies (Fig. 2A, B). Two independent RNAi constructs (v104150 and v40871 from the Vienna Drosophila Resource Center, hereinafter referred to as *KrIR#1* and *KrIR#2*, respectively) exhibited a comparable phenotype, confirming that this effect was specific to Kr depletion rather than an off-target effect (Fig. 2A, B).

To further validate the link between Kr and the adult neurogenic phenotype, we examined various *Kr* mutant alleles. Notably, while Kr null alleles were embryonic lethal, the *Irregular facet* (*Kr^If-1^*) mutation, a dominant mutation in the *Kr* locus known to cause Kr misexpression in various tissues, including the eye imaginal discs (40, 41), also exhibited EdU incorporation in multiple clones in adult brains, although at a somewhat lower frequency than *Kr* RNAi (Fig. 2B, C). These findings suggest that both reduced Kr activity and aberrant Kr expression can lead to persistence of proliferative cells in the adult central brain.

To determine the identity of these proliferative cells, we analysed the expression of NB markers. Each EdU-positive clone typically contained one to two pH3-positive cells, consistent with the presence of actively dividing cells within these clusters. As expected, these mitotic cells expressed *insc-*Gal4-driven mCD8::GFP (*insc*>mCD8::GFP) and the NB marker Miranda (Mira), confirming their NB identity (Fig. 2C, D). Notably, these mitotic NBs exhibited a crescent-like asymmetric localisation of Mira at the cell cortex during mitosis, with smaller surrounding cells lacking NB markers (Fig. 2D, G). In addition, some of the NB progeny expressed the pan-neuronal marker Elav (Fig. 2E). These observations demonstrate that active NBs are present and undergo conventional asymmetric cell divisions, producing a self-renewing NB and differentiating neuronal progeny. Intriguingly, in both *insc>KrIR* and *Kr^If-1^*mutant adult brains, we detected EdU-incorporated clones beyond 21 days after eclosion, well beyond the previously described early-adult period during which rare residual cell-cycle or proliferative activity has been reported in the adult brain (7, 46) (Fig. 2B). These findings indicate that the observed phenotype does not simply reflect a transient delay in developmental NB elimination but instead represents long-term persistence of NBs capable of sustained proliferation and neurogenesis throughout adulthood.

We next sought to determine the lineage identity of these persistent NBs. Importantly, the number and location of these NBs are reminiscent of MBNBs, which are normally eliminated during late pupal stages via apoptosis (9, 26, 31). In *insc>KrIR* or *Kr^If-1^*mutant adult brains, we consistently observed one to four EdU-positive clones per hemisphere, located on the dorsoposterior surface, matching the typical positioning of MBNBs during development (9) (Fig. 2A). To confirm their MBNB identity, we used *mb247*-Gal4, which labels MB neurons, and found that the mitotic NBs resided within the MB cell body region (Fig. 2F, G). We also used *OK107*-Gal4, which visualises the MB lineage, including MBNBs, although MBNB labelling is weaker and more variable than with the NB-specific *insc*-Gal4 driver. Using this marker, we found that a subset of EdU-positive clones co-expressed *OK107*>GFP and Mira (Fig. 2H). Together, these data identify the persistent proliferative cells as MBNBs.

### Kr functions during the pupal stage to control MBNB cell cycle exit and elimination

MBNBs are normally eliminated during late pupal development, later than other central brain NBs (9, 16, 31). To determine whether the MBNBs observed in *insc>KrIR* or *Kr^If-1^* adult brains represent persistent MBNBs that escaped developmental elimination rather than newly generated NBs, we quantified MBNB number during pupal development. In controls, we observed a gradual reduction of MBNB number during the pupal stage, whereas in *insc>KrIR* or *Kr^If-1^* mutant brains, MBNB number was maintained throughout pupal development (Fig. 3A). It is known that, preceding their elimination, MBNBs arrest their growth and gradually shrink (16). We therefore measured MBNB size and found that, in control, MBNB size was reduced in the late pupal stage compared to the early stage, whereas it remained unchanged under Kr knockdown conditions (Fig. 3B). These results indicate that Kr regulates growth arrest and elimination of MBNBs during late pupal stages.

**Figure 3.**
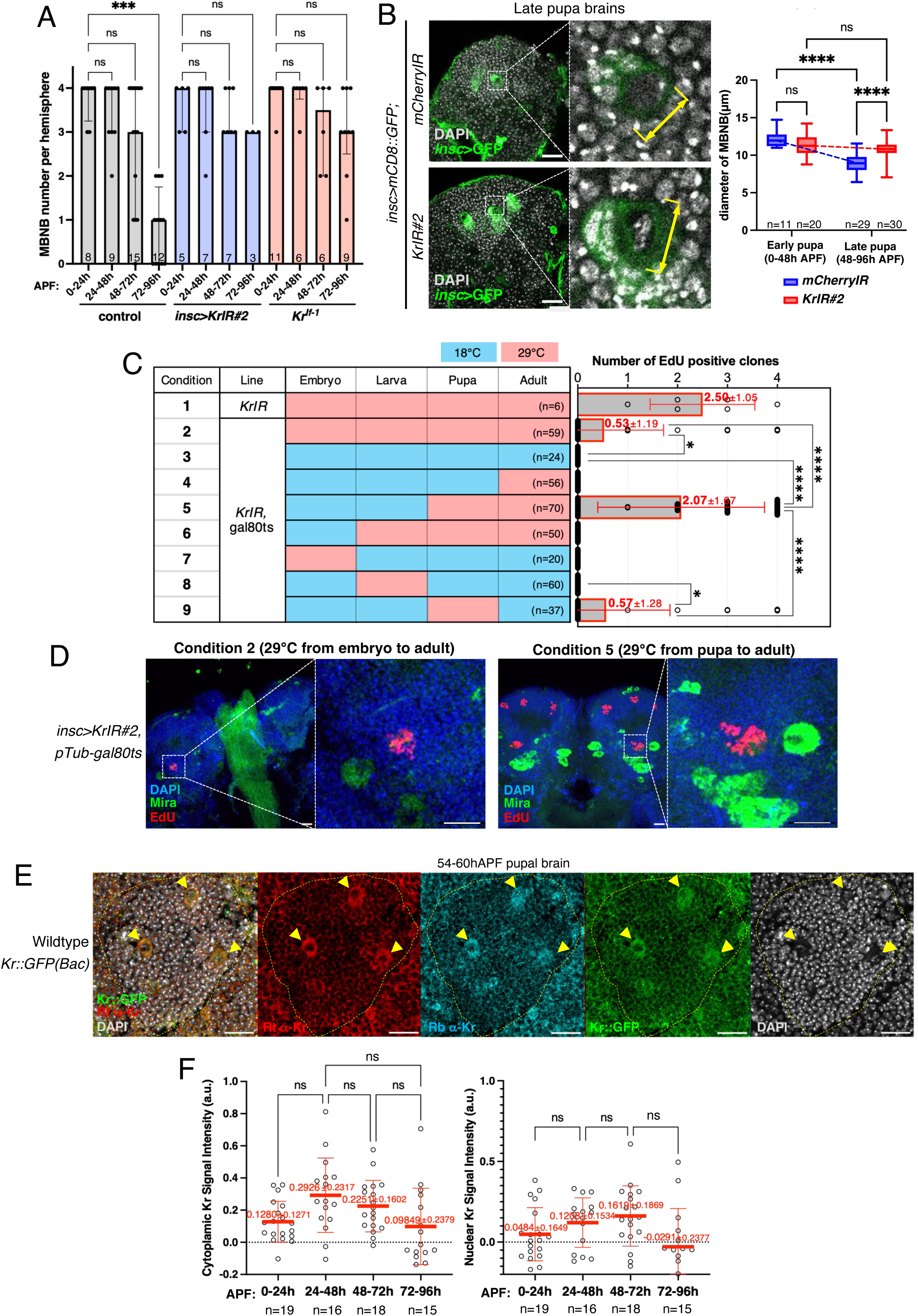
Kr is expressed in MBNBs and regulates their elimination during the pupal stage. **(A)** Quantification of MBNB number during pupal development in control, *insc>KrIR* and *Kr^If-1^* mutant brains. MBNB number progressively decreased during pupal development in controls but was maintained in *insc>KrIR* and *Kr^If-1^* mutant brains. Bars indicate medians and interquartile ranges. Numbers indicate the number of brain hemispheres analysed. Statistical significance was determined using a Kruskal–Wallis test followed by Dunn’s multiple comparisons test. **(B)** Representative confocal images and quantification of MBNB size during pupal development. MBNBs were identified by *insc*>GFP and DAPI staining. Magnified views show individual MBNBs, with yellow lines indicating the measured MBNB diameter. Scale bars: 20 µm. Quantification shows MBNB diameter in early and late pupal stages. In controls, MBNB size decreased from early to late pupal stages, whereas MBNB size was maintained upon Kr depletion. Numbers indicate the number of MBNBs analysed. Early and late pupal stages were defined based on developmental morphology, as described in Materials and methods. **(C)** Developmental stage-specific knockdown of Kr using *insc*-Gal4, *pTub*-Gal80ts and *KrIR#2*. Left: experimental scheme showing nine temperature-shift conditions. Flies maintained at 29°C (pale red boxes) induced *Kr* RNAi, whereas flies maintained at 19°C (pale blue boxes) suppressed RNAi through Gal80ts. n indicates the number of brain hemispheres analysed for EdU-positive clone counting. Right: quantification of EdU-positive clones in adult brains from each condition. Scatter dot plots show the number of EdU-positive clones per hemisphere. Greyed boxes and red bars indicate means and SDs, respectively, with values annotated above. Statistical significance was determined using pairwise Mann-Whitney U tests. **(D)** Representative confocal images of posterior adult brain regions from condition 2 and condition 5 in (C), stained for DAPI (blue), Mira (green) and EdU (red). Persistent EdU-positive clones were observed when *Kr* RNAi was induced during the pupal stage. Scale bars: 100 µm. **(E)** Representative confocal images of the MB cell body region in wild-type pupal brains at 54–60 h APF carrying the Kr::GFP(Bac) reporter. Brains were stained with rat anti-Kr antibody (Rt α-Kr, red), rabbit anti-Kr antibody (Rb α-Kr, cyan) and DAPI (grey). Kr::GFP is shown in green. MBNBs, identified by their position and large cell size within the MB cell body region, showed weak Kr signals detected by both antibodies and the Kr::GFP reporter. Scale bars: 20 µm. **(F)** Quantification of Kr signal intensity in individual MBNBs during pupal development. Scatter dot plots show normalised cytoplasmic and nuclear Kr signal intensities at different pupal stages. Thick and thin red bars indicate means and SDs, respectively. n indicates the number of brain hemispheres analysed per condition. Statistical significance was assessed separately for cytoplasmic and nuclear signals using Welch’s ANOVA followed by Dunnett T3 multiple comparisons test. No pairwise comparisons reached statistical significance after multiple-comparison correction. See Materials and methods for details of quantification. Statistical significance is indicated as follows: *p ≤ 0.05, **p ≤ 0.01, ***p ≤ 0.001, ****p < 0.0001; ns, not significant.

Kr is a well-established temporal transcription factor (tTF) in embryonic Type I NBs, where it regulates neural temporal identity together with NB proliferation and lifespan (11, 21, 24). Manipulations of embryonic tTF programmes can prolong NB proliferation, and forced expression of Kr together with additional early NB transcription factors is sufficient to induce persistent neurogenesis beyond normal developmental stages (25). However, unlike Type I NBs, MBNBs have not been shown to undergo the canonical tTF cascade (30, 57, 58). Whether Kr functions through its classical tTF role or instead has a distinct postembryonic function in MBNBs remains unclear.

To determine whether MBNB retention results from an abrogation of the NB tTF function of Kr, we first sought to determine during which developmental time windows Kr regulates MBNB elimination. Using the Gal4-Gal80ts system (59), we temporally controlled KrIR induction at different stages of development (Fig. 3C, D). We found that KrIR induction specifically during the pupal stage was sufficient to induce MBNB retention in adult brains (Conditions 5 and 9 in Fig. 3C, D), whereas induction restricted to earlier embryonic/larval stages or later adult stages had little or no effect (Conditions 4, 7 and 8). This suggests that Kr plays a critical role during the pupal stage to ensure timely cell cycle exit and subsequent elimination of MBNBs.

To assess Kr expression in MBNBs during development, we used two previously validated Kr antibodies (60, 61), together with a functional Kr::GFP(Bac) reporter line designed to report endogenous Kr expression (62), which rescued the viability of a *Kr* null mutant (*Kr^1^/Df(2R)Kr10*), supporting its functionality. We first analysed Kr expression during mid-to-late embryogenesis, and observed Kr expression in thoracic NBs, consistent with previous reports (Fig. S3A, B). However, in the presumptive MB region, where MBNBs reside, Kr was absent from MBNBs, although it was occasionally detected in *OK107*>GFP-positive neurons (Fig. S3A, B), indicating the absence of Kr in MBNBs during embryonic stages.

We next examined Kr expression during postembryonic development. In third-instar larval brains, Kr was weakly detected in MBNBs using Kr antibodies (Fig. S3C). The Kr::GFP(Bac) reporter also exhibited stronger Kr::GFP signals in single cells adjacent to MBNBs, which likely represent GMCs or immature neurons that have inherited Kr::GFP from MBNBs (Fig. S3D). In addition, strong nuclear Kr signals were also specifically observed in a few neuronal cells attached to the calyx (Fig. S3E).

In pupal brains where Kr may regulate MBNB elimination, weak Kr signals, somewhat enriched in the cytoplasm, were detected in MBNBs, as shown by both Kr antibodies and Kr::GFP(Bac) reporter (Fig. 3E). The weak Kr signals in MBNBs were observed throughout pupal development, with only modest variation between stages (Fig. 3F). These Kr signals in MBNBs were reduced in *insc>KrIR* conditions, confirming the specificity of these signals to Kr (Fig. S3F).

To independently assess Kr expression in MBNBs, we analysed publicly available transcriptomic datasets. Although tissue-level datasets (FlyAtlas2 and modENCODE) detect Kr expression in the larval and adult central nervous system, they do not resolve NB-specific expression. We therefore examined lineage-specific RNA-seq data from purified MBNBs (27), which revealed persistent but relatively low Kr transcript levels throughout larval and pupal development (Fig. S5A). These transcriptomic data are consistent with our immunostaining results, supporting the conclusion that Kr is continuously expressed at low levels in postembryonic MBNBs.

We also analysed Kr expression in MBNBs in the *Kr^If-1^* background. When we used TSA fluorescence enhancement (see Methods), we were able to detect ectopic expression of endogenous Kr, enriched in the nucleus, in MBNBs, as well as their progeny, in both *Kr^If-1^* mutant adult and pupal brains (Fig. S3G, H). These results indicate that not only reduction of Kr, but also its misexpression can induce MBNB retention, suggesting that proper regulation of Kr expression levels is required to promote timely MBNB elimination during pupal stages.

Together, these findings demonstrate that Kr functions specifically during pupal development to promote MBNB cell cycle exit and elimination, independently of its previously characterised embryonic tTF function. The persistent but low-level expression of Kr throughout postembryonic development suggests that it acts as a MBNB lineage-specific regulator that coordinates the timely transition from proliferation to terminal differentiation and elimination.

### Kr promotes MBNB cell cycle exit and elimination by regulating the Imp-Syp transition

Having established that Kr functions during the pupal stage to promote MBNB elimination, we next investigated the underlying molecular mechanism. Among the intrinsic regulators of postembryonic NB termination, the RNA-binding proteins Imp and Syp play central roles in coordinating NB survival, cell cycle exit, and neuronal fate specification through opposing temporal gradients (26, 27, 63). In MBNBs, Imp is highly expressed during early postembryonic stages, promoting proliferation and early γ-neuronal fate. As development progresses, Imp expression declines whereas Syp expression increases, driving the transition to α’β’ and subsequently αβ neuronal fates while promoting MBNB cell cycle exit and elimination (26, 27). We therefore hypothesised that Kr regulates MBNB elimination through modulation of the Imp-Syp transition.

We first examined Imp and Syp expression in *insc>KrIR* and *Kr^If-1^* mutant adult brains. In both conditions, the Imp-expressing tissue in the MB cell body region was markedly expanded compared to controls, within which mitotic MBNBs were embedded (Fig. S4A, B). To quantify the expansion of Imp-expressing cells, we measured the relative Imp- or Syp-positive area to the MB cell body region on the adult brain surface, using Eyeless (Ey) as a MB lineage marker. This analysis confirmed a significant expansion of the Imp-positive area, as well as a reduction of the Syp-positive area, in *insc>KrIR* adult brains (Fig. 4A), consistent with persistent Imp expression and incomplete acquisition of the Syp-high state in Kr-deficient MBNBs.

**Figure 4.**
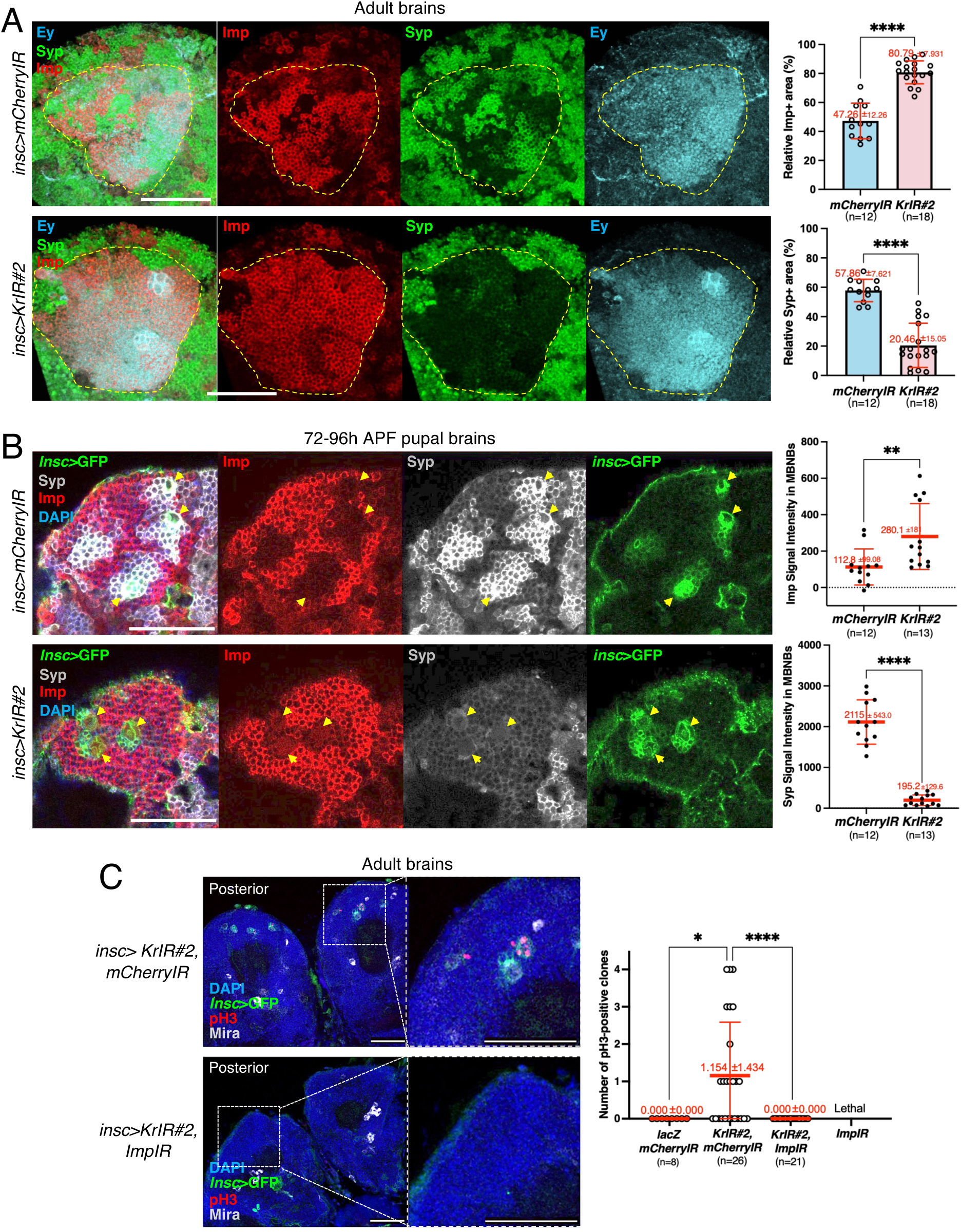
Kr promotes MBNB elimination by regulating the Imp–Syp transition. **(A)** Quantification of Imp- and Syp-expressing areas within MB cell body regions in adult brains. Left: representative 3D reconstructions of dorsoposterior adult brain surfaces from the indicated genotypes, stained for Imp (red), Syp (green) and Eyeless (Ey; blue), which marks MB lineages. Scale bars: 50 µm. Right: scatter dot plots showing the ratios of Imp-positive and Syp-positive areas to the MB cell body surface region on the dorsoposterior brain surface, quantified per hemisphere. Thick and thin red bars indicate means and SDs, respectively. n indicates the number of hemispheres analysed per condition. Statistical significance was determined using pairwise Mann–Whitney U tests. **(B)** Quantification of Imp and Syp signal intensities in MBNBs during late pupal development. Left: representative confocal images of MBNBs in 72–96 h APF pupal brains from control and *insc>KrIR#2* flies, stained for Imp (red), Syp (grey), DAPI (blue) and *insc>GFP* (green). Yellow arrowheads indicate MBNBs. Scale bars: 50 µm. Right: scatter dot plots showing normalised Imp and Syp fluorescence intensities in individual MBNBs. Thick and thin red bars indicate means and SDs, respectively. n indicates the number of MBNBs analysed. Statistical significance was determined using unpaired two-tailed t-tests; Welch’s correction was applied for Syp because variances were unequal. **(C)** Imp depletion suppresses MBNB persistence in *Kr* RNAi brains. Left: representative confocal images of adult brains from *insc>KrIR#2, mCherryIR* and *insc>KrIR#2, ImpIR*, stained for pH3 (red), insc>GFP (green), Mira (grey) and DAPI (blue). Mitotic MBNBs persisted in *insc>KrIR#2, mCherryIR* brains but were not detected in *insc>KrIR#2, ImpIR* brains. Scale bars: 100 µm. Right: scatter dot plots showing the number of mitotic MBNBs per brain. Thick and thin red bars indicate means and SDs, respectively. n indicates the number of hemispheres analysed. Statistical significance was determined using a Mann–Whitney U test. *insc>ImpIR* alone caused lethality, preventing adult brain analysis. Statistical significance is indicated as follows: *p ≤ 0.05, **p ≤ 0.01, ***p ≤ 0.001, ****p < 0.0001; ns, not significant.

Because Kr functions during pupal stages to regulate MBNB elimination, we next examined whether Kr influences the Imp-Syp transition during this period. We measured Imp and Syp expression levels in individual MBNBs during late pupal stages (72–96 h APF) and found that *insc>KrIR* MBNBs exhibited significantly higher Imp levels and lower Syp levels than controls (Fig. 4B).

To determine whether persistent Imp expression is responsible for MBNB persistence following Kr depletion, we performed co-depletion experiments by expressing both *Kr*RNAi and *Imp*RNAi in NBs. While Imp depletion alone (*insc>ImpIR*) resulted in larval lethality, co-depletion of Kr and Imp (*insc>ImpIR*, *KrIR*) partially rescued this lethality, allowing a small proportion of flies to survive to adulthood. However, these flies died within a few days of eclosion, preventing long-term EdU incorporation assays. To overcome this limitation, we assessed the presence of mitotic MBNBs using pH3 immunostaining in surviving young adult flies. Notably, while we observed mitotic MBNBs in *Kr*RNAi adult brains (*insc>KrIR, mCherryIR*), Imp co-depletion completely suppressed this phenotype (*insc>KrIR, ImpIR*; Fig. 4C, S4C). These results provide strong genetic evidence that persistent Imp expression is required for the retention and continued proliferation of MBNBs following Kr depletion.

Together, these findings demonstrate that Kr promotes the Imp-to-Syp transition during pupal development, thereby ensuring timely MBNB cell cycle exit and elimination.

### Kr and Kr-h1 function antagonistically to regulate E93 expression and MBNB termination

We next sought to determine how Kr regulates Imp expression and MBNB fate. Although Imp is required for MBNB maintenance, it is broadly expressed in many NB populations and therefore cannot readily explain the lineage-specific role of Kr in promoting MBNB cell cycle exit (27). We therefore hypothesised that Kr might regulate additional factors with more specific roles in MBNB termination.

One promising candidate is Kr-h1, another KLF family transcription factor implicated in neural development, including MB development (38, 39). Kr-h1 is a well-established target of juvenile hormone and ecdysone signalling and is known to counteract E93, an ecdysone-induced transcription factor (36, 64). Notably, E93 has been shown to be required for timely elimination of MBNBs through regulation of autophagy (31). Consistent with this relationship, MB lineage-specific transcriptomic data show that Kr-h1 expression declines sharply after the larval-to-pupal transition, coinciding with upregulation of E93 (27) (Fig. S5A). These observations are consistent with the idea that downregulation of Kr-h1 facilitates E93 activation during MBNB termination. Interestingly, Kr-h1 mutations were previously identified as enhancers of the *Kr^If-1^* eye phenotype, suggesting a functional interaction between Kr and Kr-h1 (65). Together, these findings led us to hypothesise that Kr promotes MBNB elimination, at least in part, through mechanisms that functionally oppose Kr-h1 activity and support E93-dependent termination of the MBNB lineage.

We first asked whether the antagonistic relationship between Kr-h1 and E93 also operates in the MBNB lineage. We either depleted or over-expressed Kr-h1 (*Kr-h1::FLAG*) using *insc-*Gal4 and analysed E93 expression levels by immunostaining in MBNB lineages in late pupal brains. In controls, E93 was only weakly detectable in MBNBs themselves but accumulated strongly in the surrounding progeny within the MBNB lineage (Fig. S5B). Following Kr-h1 depletion by RNAi, E93 expression was markedly increased in MBNB lineages, whereas it decreased upon Kr-h1::FLAG overexpression in NBs (Fig. S5B), confirming that the Kr-h1–E93 antagonism is conserved in the MBNB lineage.

We then investigated the functional interaction between the two KLF family proteins, Kr and Kr-h1, in the control of MBNB termination. Kr-h1 knockdown alone *(insc>Kr-h1IR*) did not prevent MBNB elimination, consistent with previous findings that Kr-h1 is largely dispensable for MB development (38, 39) (Fig. 5A). However, when Kr-h1 was co-depleted with Kr (*insc>KrIR, Kr-h1IR*), the number of retained MBNBs in adult brains was significantly reduced compared to depletion of Kr alone (*insc>KrIR, mCherryIR*), although some brains retained fewer MBNBs (Fig. 5A). These results indicate that Kr-h1 contributes to MBNB persistence following Kr depletion, supporting an antagonistic interaction between Kr and Kr-h1 in the regulation of MBNB elimination.

**Figure 5.**
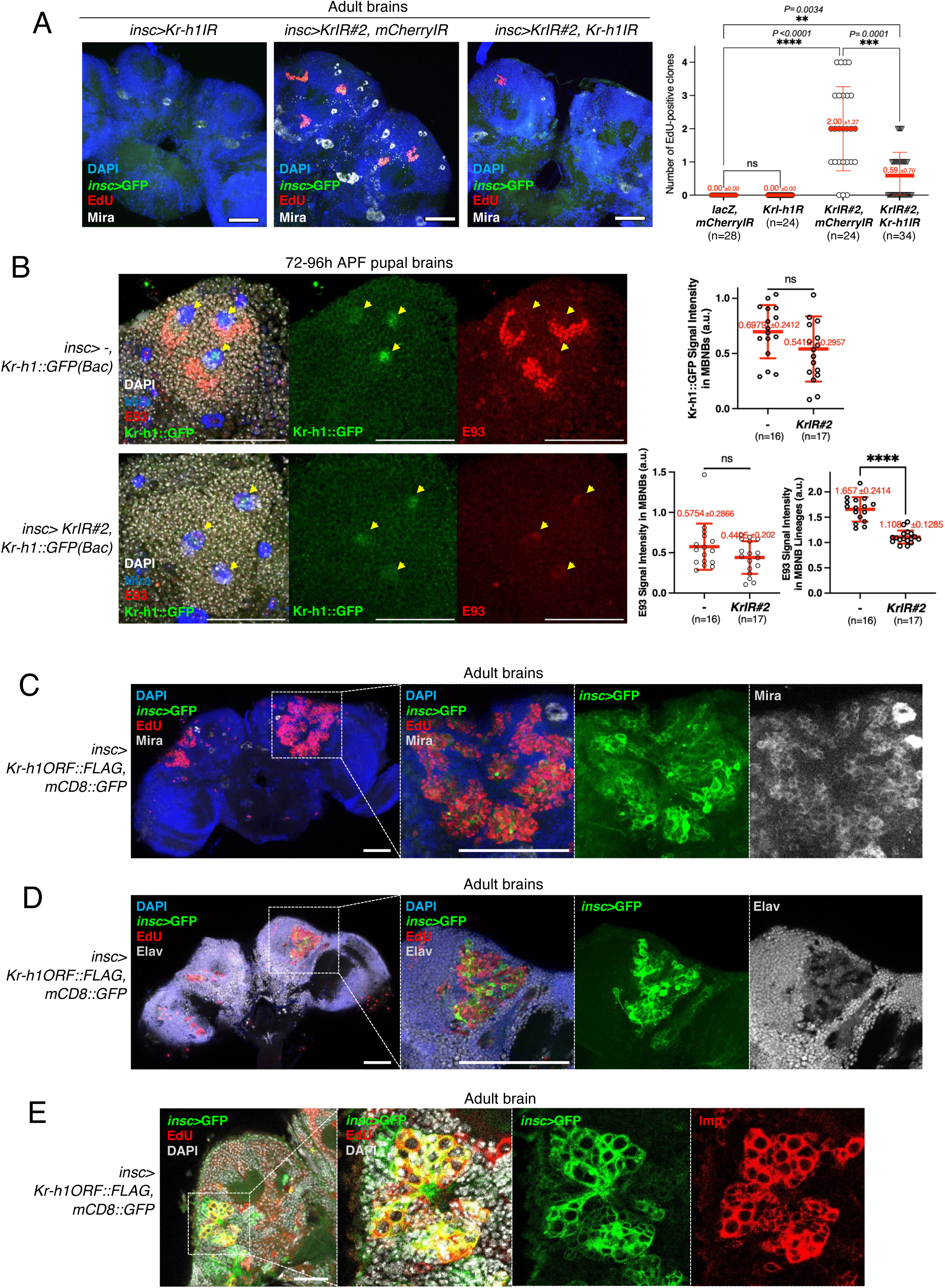
Kr antagonises Kr-h1 and supports E93 expression to promote MBNB termination. **(A)** Kr-h1 co-depletion suppresses MBNB persistence in *Kr* RNAi adult brains. Left: representative confocal images of *insc>Kr-h1IR*, *insc>KrIR#2, mCherryIR* and *insc>KrIR#2, Kr-h1IR* adult brains stained for EdU (red), *insc*>GFP (green), Mira (grey) and DAPI (blue). *Kr-h1* RNAi alone did not prevent MBNB elimination, whereas the number of persistent EdU-positive MBNB clones observed in *insc>KrIR#2, mCherryIR* brains was significantly reduced when Kr-h1 was co-depleted with Kr. Scale bars: 100 µm. Right: quantification of EdU-positive MBNB clones per hemisphere in the indicated genotypes. Scatter dot plots show individual data points, with means and SDs represented by thick and thin red bars, respectively. n indicates the number of hemispheres analysed. Statistical significance was determined using a Kruskal–Wallis test followed by Dunn’s multiple comparisons test. **(B)** Kr depletion reduces E93 expression in MBNB progeny without detectably altering Kr-h1::GFP levels in MBNBs. Left: representative confocal images of 72–96 h APF pupal brains from control *insc>Kr-h1::GFP* and Kr-depleted *insc>KrIR#2; Kr-h1::GFP* flies, stained for E93 (red), Kr-h1::GFP (green), DAPI (grey) and Mira (blue). MBNBs were identified by Mira staining and cell morphology, and the surrounding progeny region was analysed separately. Scale bars: 50 µm. Right: quantification of normalised Kr-h1::GFP signal intensity in MBNBs, and normalised E93 signal intensities in MBNBs and surrounding progeny. Scatter dot plots show individual data points, with means and SDs represented by thick and thin red bars, respectively. n indicates the number of hemispheres analysed. Statistical significance was determined using unpaired two-tailed t-tests. **(C-E)** Kr-h1 overexpression promotes NB proliferation and blocks neuronal differentiation, leading to a tumour-like NB overgrowth phenotype. (C) Representative posterior views of adult brains overexpressing Kr-h1 in NBs (*insc>Kr-h1ORF::FLAG*), stained for DAPI (blue), insc>GFP (green), EdU (red) and Mira (grey), with magnified views of the MB cell body regions shown on the right. Most EdU-positive cells within enlarged clones co-expressed Mira and *insc*>GFP, indicating their NB-like status and a failure in differentiation. Scale bars: 100 µm. (D) Confocal images of *insc>Kr-h1ORF::FLAG* adult brains stained for DAPI (blue), *insc*>GFP (green), EdU (red) and Elav (grey). EdU-labelled clones in Kr-h1-overexpressing brains contained very few Elav-positive neurons, indicating impaired neuronal differentiation. Scale bars: 100 µm. (E) Confocal images of *insc>Kr-h1ORF::FLAG* adult brains stained for DAPI (grey), *insc*>GFP (green) and Imp (red). Scale bars: 20 µm. Statistical significance is indicated as follows: *p ≤ 0.05, **p ≤ 0.01, ***p ≤ 0.001, ****p < 0.0001; ns, not significant.

We next investigated whether Kr regulates Kr-h1 expression in MBNBs. Kr-h1 was visualised using a functional Kr-h1::GFP(Bac) reporter. As expected from the MBNB lineage-specific transcriptomic data (Fig. S5A), Kr-h1::GFP showed clear nuclear signals in MBNBs during third instar larval stage but became barely detectable during late pupal development (Fig. 5B, S5C). Kr depletion did not significantly alter Kr-h1::GFP levels in MBNBs. Likewise, the weak E93 signal detectable in MBNBs themselves remained largely unchanged. In contrast, Kr depletion almost completely abolished E93 expression in the surrounding progeny of the MBNB lineage (Fig. 5B). These findings suggest that Kr is required for proper activation of E93 in MBNB lineages independently of detectable changes in Kr-h1 expression.

To further examine Kr-h1’s function, we overexpressed Kr-h1 in MBNBs using *insc*-Gal4. Strikingly, Kr-h1::FLAG induction led to a dramatic increase in the number of proliferating NBs, including MBNBs, which continued Imp expression, in the adult central brain, a phenotype markedly more severe than that observed in Kr depletion (Fig. 5C-E). These data indicate that Kr-h1 promotes NB proliferation and prevents their elimination. Notably, beyond its effect on NB persistence, Kr-h1 overexpression appears to also disrupt normal asymmetric division and differentiation of NBs. In *insc>Kr-h1::FLAG* adult brains, most EdU-labelled cells retained expression of the NB markers Mira and *insc*>GFP, while very few cells expressed Elav, suggesting a failure in neuronal differentiation (Fig. 5C, D). This phenotype resembles NB tumour-like overgrowths observed in *brat, numb* or *pros* mutant NBs, in which progeny fail to differentiate and instead retain a progenitor-like state (66).

Together, these findings support a model in which Kr and Kr-h1 function antagonistically to regulate MBNB termination. Kr promotes the Imp-to-Syp transition and supports E93 activation within MBNB lineages, thereby driving MBNB cell cycle exit and elimination, whereas Kr-h1 counteracts this programme by repressing E93, thereby promoting continued NB proliferation and self-renewal.

### Kr coordinates MBNB termination and mushroom body development

Our findings indicate that Kr promotes MBNB cell cycle exit by promoting the Imp-to-Syp transition and supporting E93 activation within the MBNB lineage during late pupal development. Given the established role of Imp in regulating temporal transitions of MB neuronal identity, we hypothesised that Kr may influence MB development beyond its role in NB elimination.

To test this, we examined MB morphology in *insc>KrIR* and *Kr^If-1^* adult brains. These flies developed normally, and *OK107*>GFP-labelled MB structures showed no significant abnormalities in their overall architecture (Fig. 6A). Despite largely normal overall MB morphology, FasII staining revealed subtle abnormalities in the α/β lobes following Kr depletion, with the lobes appearing thinner and more curved than in controls (Fig. 6B). These abnormalities are consistent with defective or delayed α/β lobe formation during MB development.

**Figure 6.**
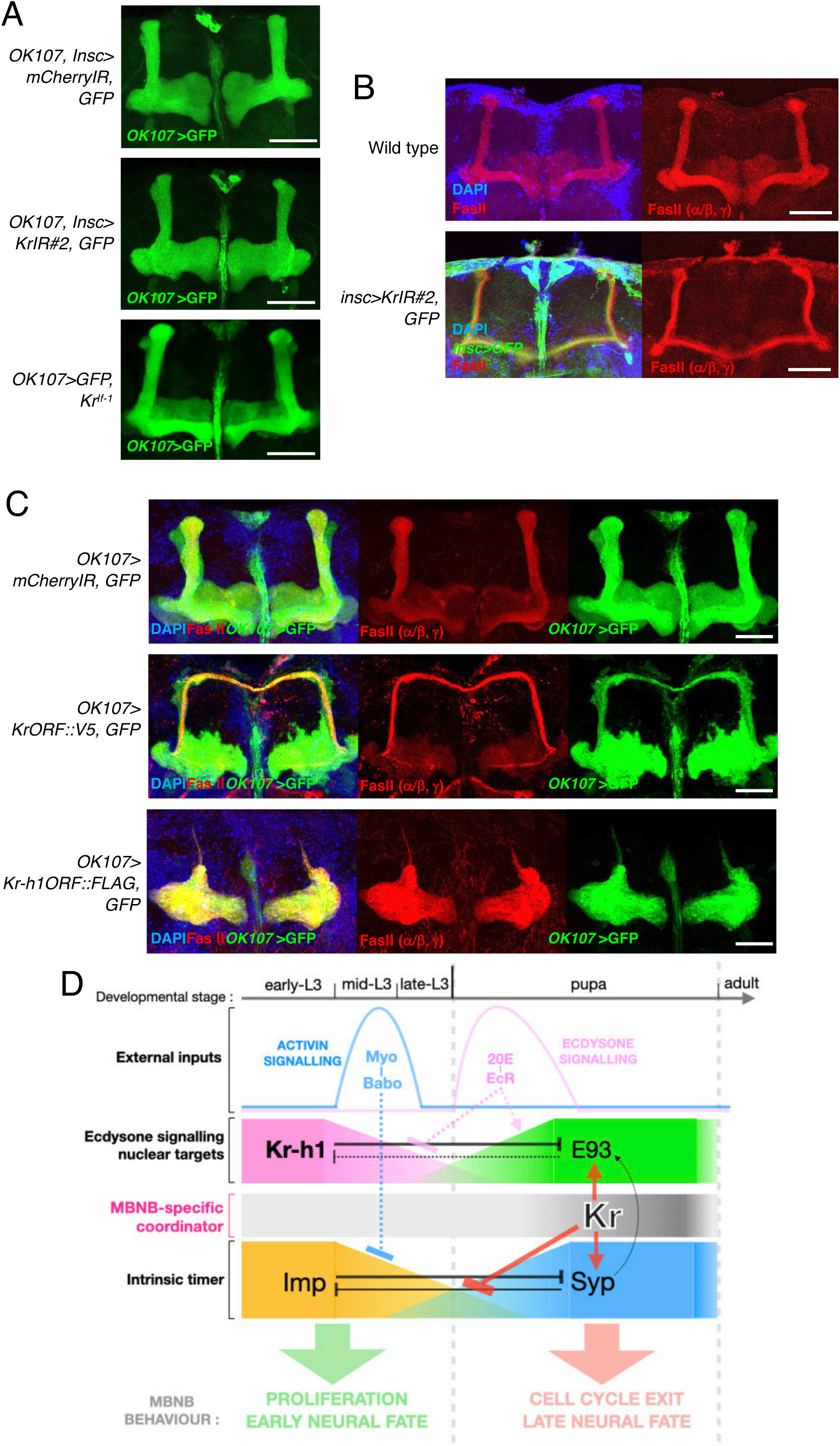
Kr coordinates MBNB termination and mushroom body development. **(A)** Representative confocal images of anterior views of adult brains from the indicated genotypes, with MB lobe structures visualised by *OK107*>GFP (green). No overt structural abnormalities were observed in MB lobes following Kr depletion or in *Kr^If-1^* mutant adult brains. Scale bars: 50 µm. **(B)** Representative images of MB α/β lobe morphology in control and *insc>KrIR#2* adult brains, visualised by Fasciclin II (FasII) staining (red). Kr depletion led to thinner and more curved α/β lobes compared to controls. Scale bars: 50 μm. **(C)** Effects of Kr or Kr-h1 overexpression in MB lineages on MB development. Representative confocal images of anterior views of adult brains from *OK107>mCherryIR* control, *OK107>KrIR#2*, *OK107>KrORF::V5* and *OK107>Kr-h1ORF::FLAG* flies, are shown, with MBs visualised by *OK107*>GFP (green) and FasII staining (red). Kr overexpression caused high lethality; surviving *OK107>KrORF::V5* adult flies exhibited severe disorganisation of MB lobes, including pronounced reduction of α/β and α′/β′ lobes and extensive disorganisation of γ lobes. Kr-h1 overexpression also caused severe MB disorganisation, with persistent FasII-positive structures in the β and/or β′ lobe regions. Scale bars: 50 μm. **(D)** Proposed model for Kr function in coordinating MBNB termination. During larval development, high Imp and Kr-h1 levels maintain MBNB proliferation and early neuronal fate. As development progresses into the pupal stage, the intrinsic temporal programme shifts towards high Syp and E93 expression, promoting cell cycle exit, neuronal maturation and eventual MBNB elimination. Kr acts as an MBNB-specific coordinator that promotes this transition by suppressing the persistent Imp-dominated programme while facilitating Syp and E93 activation within the MBNB lineage. In parallel, Kr-h1 antagonises E93 as part of the ecdysone-responsive transcriptional programme. Solid regulatory arrows indicate relationships supported by data in this study, whereas dashed inhibitory arrows indicate regulatory interactions reported in other developmental contexts or inferred from the literature but not directly tested in MBNBs in this study. Activin/Babo signalling and ecdysone signalling provide extrinsic developmental cues that regulate the timing of this transition. Our data suggest that Kr does not primarily regulate E93 through detectable changes in Kr-h1 expression, but instead promotes the E93-dependent termination programme through a parallel pathway while simultaneously coordinating the Imp–Syp temporal transition. Loss of Kr disrupts these coordinated transitions, resulting in prolonged MBNB proliferation and continued neurogenesis in adult brains.

To further explore the role of Kr and Kr-h1 in MB development, we examined the effects of their overexpression. Overexpression of Kr (KrORF::V5) using *insc*-Gal4 resulted in early larval lethality, precluding further analysis. We therefore used *OK107*-Gal4, which is expressed more weakly in MBNBs than *insc*-Gal4, to drive Kr or Kr-h1 overexpression within the MB lineage. Although most *OK107*>Kr::V5 animals also failed to survive, a small proportion reached adulthood and were analysed. Unlike the largely normal MB morphology observed in *insc>KrIR* and *Kr^If-1^* adult brains, these surviving *OK107>KrORF::V5* flies exhibited severe morphological MB abnormalities, with marked reduction of the α/β and α’/β’ lobes together with extensive disorganisation of the γ lobes (Fig. 6C). *OK107>Kr-h1ORF::FLAG* flies also exhibited similarly disrupted MB morphology. Interestingly, strong FasII immunoreactivity remained associated with residual structures in the β/β′ lobe region (Fig. 6C). Together, these findings indicate that appropriate regulation of both Kr and Kr-h1 activity is required not only for MBNB termination but also for normal MB morphogenesis.

## Discussion

### The Adult *Drosophila* Central Brain Is Predominantly Postmitotic but Retains Latent Neurogenic Capacity

Consistent with previous studies (9, 16, 17), our analysis confirms that the adult *Drosophila* central brain is predominantly postmitotic under physiological conditions. In control adult brains, we detected virtually no mitotic figures or EdU incorporation, and nearly all cells displayed a G1/G0-like Fly-FUCCI profile (Fig. 1A and Fig. 2A, B). It should be noted, however, that our results do not exclude the possibility that largely quiescent or extremely slow-dividing latent progenitor-like cells exist in the adult brain and can be reactivated under specific conditions, such as injury or excessive neuronal loss (42-44). Our transcriptomic analysis of CCR expression using the existing adult and larval brain scRNA-seq datasets (47, 48) further supports a robustly postmitotic state of the adult central brain (Fig. 1B, C and Fig. S1). Most positive CCRs, including cyclins and Cdks, were broadly suppressed in adult neuronal and glial clusters, whereas several negative CCRs, such as *fzr*, *grp* and *geminin*, remained expressed across neural cell types. These observations suggest that coordinated repression of positive CCRs, rather than a simple upregulation of inhibitory factors, is likely to be a key feature of the postmitotic state in adult brain tissue, which may be important for preventing unintended cell cycle re-entry and preserving neural integrity.

Consistent with this idea, forced expression of E2F1-Dp and Cdk2-CycE was sufficient to induce mitotic entry in *TH*-Gal4-positive dopaminergic neurons and *mb247*-Gal4-positive mushroom body neurons (Fig. S2). This ectopic mitotic entry was associated with Dcp-1 staining and reduced adult survival when induced in dopaminergic neurons using *TH*-Gal4 (Fig. S2B, C). Aberrant cell cycle entry in postmitotic neurons has been linked to neuronal apoptosis and neurodegeneration in multiple contexts, including E2F/Rb-dependent responses to DNA damage (50, 67, 68). Thus, our data indicate that forced activation of positive CCRs in postmitotic neurons can induce aberrant cell-cycle re-entry, which may trigger cell death and compromise adult survival. However, the precise causes of neuronal death and reduced survival in our system remain to be determined, and we cannot exclude contributions from *TH-*Gal4 activity outside dopaminergic neurons or other indirect effects of the manipulation.

### Kr Functions in the Timely Elimination of MBNBs to Prevent Adult Neurogenesis

A central finding of this study is that Kr has a previously unrecognised postembryonic function in controlling the timely termination of MBNBs (Fig. 2 and Fig. 3). Kr is best known as a gap gene and as an early tTF in embryonic Type I NBs, where it shows a transient burst of expression and contributes to temporal identity specification, NB proliferation and NB lifespan (11, 21, 24, 25). However, Kr does not appear to act as a conventional embryonic tTF in MBNBs. Kr was not detected in embryonic MBNBs at levels comparable to those observed in thoracic NBs, whereas it was expressed at low and relatively constant levels in MBNBs throughout postembryonic development (Fig. 3E, F and Fig. S3). MBNBs persist until late pupal stages, much longer than most other central brain NBs, before undergoing cell cycle exit and elimination (9, 16), and Kr supports this MBNB-specific termination programme. Pupal-stage-specific Kr depletion was sufficient to cause MBNB retention in adult brains, whereas knockdown during earlier embryonic or larval stages did not produce the same effect (Fig. 3C, D). Importantly, this Kr function is highly lineage-specific: upon NB-specific Kr depletion, MBNBs persist and continue to divide in the adult brain, while other central brain NBs are still eliminated before adulthood. Thus, Kr acts as an MBNB lineage-specific factor that controls their unique cell cycle and termination programme. Notably, proliferative MBNBs were detected even more than three weeks after eclosion, indicating that Kr controls long-term MBNB behaviour and neurogenic potential, rather than simply determining their elimination timing.

The analysis of the classic *Irregular facet* mutation further supports the importance of precise regulation of Kr expression. The *Kr^If-1^* allele, originally identified through its adult eye phenotype and known to misregulate Kr expression (40, 41), also caused ectopic Kr expression in MBNBs during late pupal development and led to their persistence in adult brains (Fig. 2 and Fig. S3G). Thus, normal MBNB termination requires tight control of the timing and level of Kr activity.

Together, these findings highlight MBNBs as an unusually flexible neurogenic population and suggest that relatively subtle changes in transcriptional regulation can unlock prolonged neurogenesis in an otherwise postmitotic adult brain. The *Irregular facet* mutation provides one of the few examples in which a classic developmental mutation reveals latent adult neurogenic potential in the *Drosophila* brain. Notably, adult MB neurogenesis has been reported in other insects, including crickets and moths (18, 19). It will therefore be interesting to determine whether species-specific differences in the timing, level or regulation of Kr/KLF activity in MBNBs contribute to evolutionary variation in adult neurogenic capacity.

### Kr Coordinates the Imp-to-Syp Transition and E93 Activation, while Kr-h1 Opposes MBNB Termination

Mechanistically, our results place Kr upstream of the Imp-to-Syp transition in pupal MBNBs (Fig. 4, Fig. S4 and Fig. 6D). The opposing Imp-Syp gradient is a central regulator of NB temporal progression (26, 27, 63), and, in MBNBs, delayed termination is accompanied by a late transition from the Imp-high to the Syp-high state (26, 27, 30). In Kr-depleted brains, MBNBs maintained high Imp expression and failed to fully acquire Syp expression, indicating that Kr is required for the proper progression of this temporal transition (Fig. 4A, B). Importantly, co-depletion of Imp suppressed MBNB retention caused by Kr depletion, demonstrating that persistent Imp expression is required to sustain MBNB proliferation in the absence of Kr (Fig. 4C). Thus, Kr promotes MBNB termination at least in part by limiting persistent Imp expression and allowing the normal Imp-to-Syp transition.

In addition to regulating the Imp-Syp transition, Kr also supports E93 expression within the MBNB lineage (Fig. 5). E93 is of particular interest because, unlike the broadly acting Imp-Syp temporal programme in many NBs, E93 specifically promotes MBNB termination during late pupal development, partly through regulation of autophagy (31). In our experiments, Kr depletion markedly reduced E93 expression in the MBNB lineage, particularly in the neuronal progeny (Fig. 5B). These observations suggest that Kr is required for proper E93 activation during MBNB termination, although the precise cell type and mechanism through which Kr supports E93 expression remain to be determined.

Our data also reveal an antagonistic role for another KLF family transcription factor, Kr-h1, in MBNB termination (Fig. 5 and Fig. 6D). Kr-h1 is a hormone-responsive transcription factor and a well-established antagonist of E93 in broader developmental contexts (36, 37, 64). Consistent with this relationship, Kr-h1 depletion increased E93 expression in the MBNB lineage, whereas Kr-h1 overexpression strongly suppressed it (Fig. S5). Functionally, Kr-h1 knockdown partially suppressed MBNB retention caused by Kr depletion, whereas Kr-h1 overexpression caused tumour-like MBNB overproliferation accompanied by sustained Imp expression, supporting the idea that Kr-h1 opposes the termination programme promoted by Kr (Fig. 5A, C-E). However, Kr depletion did not detectably increase Kr-h1::GFP levels in late pupal MBNBs (Fig. 5B), suggesting that Kr and Kr-h1 do not act simply through a linear Kr–Kr-h1 pathway, but rather through convergent or parallel regulation of shared downstream processes. Notably, Kr-h1 overexpression caused a much more severe phenotype than Kr depletion, producing extensive overproliferation and a strong failure of neuronal differentiation in NBs beyond MBNBs (Fig. 5C-E). This suggests that Kr-h1 may have additional activity to maintain or reinforce a progenitor-like state in NBs, beyond its regulation of E93. Given that Chinmo acts downstream of the Imp/Syp temporal programme and can cooperate with Imp to maintain progenitor-like or tumour-propagating states (27, 69, 70), it will be interesting to determine whether Chinmo is also maintained in Kr-depleted MBNBs or Kr-h1-overexpressing NB-like cells.

Together, our findings support a model in which Kr coordinates multiple cell-intrinsic and extrinsic mechanisms to promote the MBNB-specific termination programme (Fig. 6D). Kr promotes the Imp-to-Syp transition and supports E93 activation within the MBNB lineage, thereby enabling timely cell cycle exit and elimination. Kr-h1 counteracts this programme, at least in part by repressing E93. This antagonistic balance between Kr and Kr-h1 may provide a mechanism by which MBNBs integrate intrinsic temporal progression with extrinsic developmental cues to control the timing of neural progenitor termination.

The exact mechanisms through which Kr coordinates these multiple pathways remain unclear. However, it is noteworthy that impaired Activin/Babo signalling in MBNBs produces outcomes related to Kr depletion, including prolonged Imp expression and disrupted MB temporal fate progression, and that Kr-h1 removal can also suppress these defects (39). Since Activin/Babo signalling promotes timely Imp downregulation in MBNBs (30), Kr may converge with Activin/Babo-regulated temporal progression to coordinate the delayed Imp-to-Syp transition and E93-dependent termination of MBNBs. Notably, the weak Kr signal in pupal MBNBs appeared partly enriched in the cytoplasm (Fig. 3E), raising the possibility that Kr may not act solely as a canonical nuclear transcription factor in this context. Future studies should determine how Kr activity is regulated in pupal MBNBs and whether it intersects with Activin/Babo signalling, E93 regulation, or additional non-canonical mechanisms that contribute to MBNB termination.

### Kr Links MBNB Termination with Mushroom Body Development

Beyond MBNB cell cycle exit and elimination, our data suggest that Kr influences MB development (Fig. 6). Kr depletion caused relatively subtle defects in MB morphology, most notably in the α/β lobes, whereas overexpression of Kr or Kr-h1 produced much more severe disruption of MB lobe architecture. These phenotypes suggest that appropriate regulation of Kr and Kr-h1 activity is important not only for MBNB termination, but also for normal MB lineage progression and morphogenesis.

These developmental defects may reflect altered temporal identity progression of MB neurons produced during development. Since the Imp-Syp gradient governs neuronal fate progression (26, 27, 30), persistent Imp expression in Kr-depleted MBNBs could bias neuronal progeny toward earlier identities or delay their transition to later identities. However, we have not directly determined the identity or connectivity of neurons produced by persistent MBNBs in Kr-depleted or *Kr^If-1^* mutant adult brains. Another important question is how this altered MB neurogenesis affects MB function and, potentially, brain function more broadly. Given the critical role of the MB in olfactory learning and memory (13, 14), future studies will be required to determine whether prolonged MB neurogenesis alters MB circuit function, learning, plasticity or ageing.

### KLFs in Neural Progenitor Termination and Latent Neurogenic Potential

Our study has uncovered previously unrecognised roles of two KLF family transcription factors, Kr and Kr-h1, in controlling the proliferation and neurogenic potential of a specific neural progenitor population. KLF family transcription factors are evolutionarily conserved regulators of development, stem cell behaviour and ageing (32, 33). In mammals, KLF4 is a well-known pluripotency factor and has been implicated in protecting neural stem cells from senescence (71, 72), whereas KLF9 regulates neural stem cell quiescence and can influence adult hippocampal neurogenesis (73, 74). Although the mechanisms and developmental contexts are clearly different, these parallels suggest that KLF proteins may have a broader role in tuning the balance between progenitor maintenance, quiescence and termination across species.

In this context, the antagonistic balance between Kr and Kr-h1 in *Drosophila* MBNBs provides a genetically tractable model for understanding the mechanisms by which progenitor lifespan and neurogenic potential are regulated in a lineage-specific manner. MBNBs are particularly informative because their termination is naturally delayed compared with most other central brain NBs, and because the timing of MBNB termination and the persistence of MB neurogenesis appear to vary across insect species. Thus, understanding how Kr/KLF-dependent programmes regulate MBNB termination may provide insight into how neurogenic potential is developmentally restricted, maintained or extended, and may help explain evolutionary variation in adult neurogenic capacity. More broadly, this study highlights how small changes in developmental transcriptional programmes can alter the boundary between a postmitotic adult brain and a brain with persistent neurogenic capacity.

### Materials and Methods Fly stocks and culture

The *Drosophila* lines used in this study are as follows. Wild-type and control stocks, *Oregon R* and *w^1118^*, and a *Kr^If-1^* mutant stock, *Kr^If-1^/Cyo; Sb/TM6B* were obtained from Cambridge Drosophila Facility. An additional *Kr^If-1^* mutant stock *w^1118^; Kr^If-1^/CyO* (BCF360) and *Kr* RNAi lines (v104150, v40871) were obtained from Shanghai Fly Center and VDRC, respectively. The Gal4 driver lines specific for the mushroom body neurons, *mb247-Gal4* (a gift from Dr Margaret Ho) and *OK107-Gal4; UAS-mCD8::GFP* (a gift from Dr Yi Zhong). The NB-specific Gal4 driver lines *, insc-Gal4* (BL8751) from the Bloomington Drosophila Stock Center (BDSC), *UAS-dicer2*; *insc*-*Gal4, UAS-mCD8::GFP*/CyO (a gift from Dr Eugen Knoblich). *pTub-gal80ts* lines (BL7107, BL7108), a GFP reporter line, *UAS-mCD8::GFP* (BL5130), a Kr overexpression line *UAS-Kr-V5* (BL83301), a loss-of-function *Kr* mutant *Kr^1^* (BL3494) and a *Kr* deficiency line, *Df(2R)Kr10* (BL4961), a Kr::GFP(Bac) reporter line *PBac[Kr-GFP.FPTB]* (BL56152), and a Kr-h1::GFP(Bac) reporter line *PBac[Kr-h1-GFP.FPTB]* (BL96786), and a UAS-Dcr2 line(24650). An *Imp* RNAi line (THU55645) from Tsinghua Fly Center. *Kr-h1* RNAi line (BL50685) and *UAS-Kr-h1::FLAG* (kind gifts from Qianyu He). A Fly-FUCCI line *pUbq-GFP-E2F1, mRFP-NLS-CycB* (a kind gift from Dr Bruce Edgar).

All *Drosophila* lines were maintained and cultured with common cornmeal agar media, with 12 hr-12 hr day-night cycles, and at 50-70% humidity. All fly experiments were conducted at 25°C except for RNAi and overexpression experiments in which flies were grown at 29°C for higher Gal4-dependent induction, as specifically noted. 20 to 30 flies were contained in each tube and flipped them into new tubes every two to three days. Fly crossing was done with around 10 virgins and 8 males with specific genotypes in each tube. In most crosses, virgins of Gal4 driver lines were collected and crossed with the male flies with UAS elements. The progeny with the specific genotypes were selected based on visible genetic markers for analysis.

### Antibodies

Primary antibodies used in this study are as follows: rabbit Phospho-Histone H3 (Ser10) antibody (Invitrogen PA5-17869, 1:400), rat anti-Elav (Developmental Studies Hybridoma Bank, DSHB, 7E8A10 1:50,), rat anti-Mira (1:400) and rat anti-Dpn (1:100, kind gifts from Dr Chris Doe), rabbit anti-Kr antibody (a gift from Dr Chris Rushlow, 1:1000)(60), rat anti-Kr antibody (from Asian Distribution Center for Segmentation Antibodies, 1:100)(61, 75), guinea pig anti-Syp (a gift from Ilan Davis, 1:200), rat anti-Imp and rabbit anti-Syp (kind gifts from Dr Claude Desplan, 1:200 and 1:1000), mouse anti-Eyeless (DSHB, AB2253542), guinea pig anti-E93 (a kind gift from Dr Chris Doe), mouse anti-FasII (DSHB, 1D4, 1:40), rabbit anti-Cleaved *Drosophila* DCP-1 (Cell Signaling Technology, 1:100), mouse anti-V5 antibody (Invitrogen, SV5-Pk1), and chicken anti-GFP (ab13970). Secondary antibodies used in this study are goat anti-mouse, anti-rat, anti-rabbit, anti-guinea pig and anti-chicken secondary antibodies conjugated with Alexa Fluor 488, 568 or 647 (Invitrogen, 1:500). DAPI was used to stain DNA (Cell Signaling Technology, 1:1000).

### Immunostaining of *Drosophila* brains and embryos

*Drosophila* adult brains were dissected in 1x PBS following the protocol described previously (76). The brains were then fixed in freshly made fixative containing 8% formaldehyde, 1x PBS, 0.5mM EGTA and 5mM MgCl_2_, at room temperature for 1 hour. The fixed brain samples were then washed in PBST (1x PBS, 0.3% Triton-X), blocked with 3% BSA in PBST (PBSTB) at room temperature for 1 hour, and then incubated in PBSTB containing appropriate primary antibodies at 4°C overnight. After washes, the samples were further incubated in PBSTB containing secondary antibodies and DAPI (Cell Signaling Technology) at room temperature for 3 hours, then washed in PBST and kept in mounting media (Vectashield, VectorLab). The brain samples were then mounted onto microscope slides and kept in 4°C till imaging.

For pupal brains, pupae were removed from the pupal case and fixed in freshly made fixative containing 8% formaldehyde, at room temperature for 40 minutes, and after washing fixed pupal brains were then carefully dissected in 1x PBS.

Larval brain-eye complexes were fixed in freshly made fixative containing 4% formaldehyde, 1x PBS, 0.5mM EGTA and 5mM MgCl_2_, at room temperature for 30 minutes. After washing, blocking, and staining steps mentioned as above, larval brains were carefully separated from the complexes and mounted onto slides.

For embryo staining, parental flies were kept in embryo collection cages with apple juice agar plates supplemented with yeast paste. Embryos were collected at the desired developmental stages using a brush, transferred to baskets, dechorionated in 50% bleach and rinsed thoroughly with water. Dechorionated embryos were transferred to glass vials containing 8% formaldehyde in PBS and heptane at a 1:1 ratio and fixed for 20 min at room temperature. After removal of the lower aqueous phase, embryos were devitellinised by adding an equal volume of −20°C methanol and shaking for 1 min. Embryos were washed three times in methanol for 5 min each and then rehydrated in PBS and PBST for 15 min each. Immunostaining was performed as described above, except that embryos were blocked in PBTA (PBS containing 0.05% Triton X-100 and 1% BSA) for 1 h at room temperature.

### Tyramide signal amplification (TSA) in *Drosophila* brains

Tyramide signal amplification (TSA), based on HRP-mediated catalysed reporter deposition, was used to amplify weak immunofluorescence signals (77, 78). *Drosophila* pupal or adult brains were dissected as described above and fixed in 4% paraformaldehyde for 25 min at room temperature. After fixation, samples were permeabilised and washed three times for 10 min each in PBS containing 1% Triton X-100 (PBST). Endogenous peroxidase activity was quenched by incubating the samples in 3% H₂O₂ for 1 h at room temperature, followed by three 10-min washes in PBST. The brains were then blocked in PBST containing 3% BSA for 30 min at room temperature and incubated with primary antibodies diluted in blocking solution overnight at 4°C. For Kr staining, the rat anti-Kr antibody was pre-absorbed before use and applied at empirically optimised dilutions. After three 10-min washes in PBST, samples were incubated with HRP-conjugated secondary antibodies, together with DAPI where appropriate, for 3 h at room temperature. The samples were then washed three times for 10 min each in PBST, and the TSA reaction was carried out for 10 min at room temperature using the Yeasen Cy3 TSA Fluorescence System Kit (Yeasen, Cat# 60406ES) with Cy3-tyramide diluted 1:500 in amplification buffer, according to the manufacturer’s instructions. After three further 10-min washes in PBST, samples were subjected to additional conventional immunostaining where required, including staining with anti-Miranda antibody followed by the appropriate fluorescent secondary antibody. Before mounting, brains were further cleaned by removing residual non-brain tissues. Samples were then mounted on microscope slides and stored at 4°C until imaging.

### Confocal imaging and image processing

Confocal images were acquired using a Nikon C2 confocal microscope with 405, 488, 561/568 and 640/647 nm laser lines, as appropriate for each experiment. Image reconstruction and analysis were performed using NIS-Elements AR 5.20 and ImageJ/Fiji. Unless otherwise stated, representative fluorescence images are shown as maximum-intensity projections of confocal z-stacks. For display purposes, brightness and contrast were adjusted uniformly across comparable images within the same experiment. In some representative images, mild image processing, including a 0.5-pixel Gaussian blur or despeckle filtering, was applied only to improve visualisation. Quantitative analyses were performed on unprocessed or equivalently processed source images, and identical analysis settings were applied to all conditions within each experiment.

For quantification of Imp- and Syp-positive areas in Fig. 4A, 3D reconstructions were generated from confocal z-stacks using the Volume View/Max Intensity Projection function in NIS-Elements. Reconstructed images were rotated so that the mushroom body cell body region on the dorsoposterior brain surface faced the viewer. The MB cell body surface area and Imp-or Syp-positive areas within this region were then measured from these consistently oriented reconstructed images, as described below.

### EdU labelling of proliferating cells in *Drosophila* adult brains

EdU labelling was performed to detect proliferating cells in *Drosophila* adult brains, following the protocol outlined by Siegrist et al. (2010) (16). EdU powder was dissolved in a 5% sucrose solution to achieve a final concentration of 0.5 mg/ml, and this solution was applied to paper to replace the regular food source in the culture vials. One- to three-day-old young adult flies were then collected and raised until the flies reached certain ages, and then fed with EdU-supplemented food for a period of three days prior to brain dissection. For EdU labelling chase experiments, EdU exposure was ceased after the initial labelling period, and the flies were subsequently maintained for an additional three, seven, or fourteen days before brain dissection.

Following dissection, adult brains were subjected to immunofluorescence staining using standard procedures, with the exception of an additional step for EdU detection. EdU Click labelling was performed in the dark for 30 minutes prior to primary antibody incubation, following the manufacturer’s instructions (APExBIO EdU Imaging Kits Cy3, #K1075). EdU-incorporated cell clones were visualised and manually counted in adult brains using a Nikon Ti2 fluorescence microscope equipped with a 20x objective lens.

### Quantification of MBNB number in pupal brains (Fig. 3A)

Control, *insc>KrIR* and *Kr^If-1^* mutant pupae were collected over a 24 h period and incubated at 25°C for 0–24, 24–48, 48–72 or 72–96 h after puparium formation (APF) before dissection. Pupal brains were fixed and immunostained as described above. MBNBs were identified based on their characteristic dorsoposterior location, rounded nuclear morphology, large cell size and proximity to the developing mushroom body calyx. The number of MBNBs per brain hemisphere was manually counted using ImageJ/Fiji.

### Quantification of MBNB diameter in pupal brains (Fig. 3B)

Pupal brains were dissected and fixed using a standard immunostaining protocol as described above. Brains from the *insc-Gal4>UAS-mCD8::GFP* line were used to visualise MBNBs. Brains were mounted with the posterior side facing the coverslip and imaged on a Nikon C2 confocal microscope. Z-stacks were captured across the depth of the brain that included the entire MB cell body area and part of the MB calyx using a 1.5 µm z-step in two channels: DAPI and GFP. The MBNB diameter was measured between the two furthest points within the circular signal defined by mCD8::GFP.

### Developmental stage-specific Kr knockdown using Gal80ts (Fig. 3C, D)

Developmental stage-specific Kr knockdown was performed using *insc-Gal4, pTub-Gal80ts* and *<u>UAS-KrIR</u>*. Crosses were maintained under temperature-shift conditions to restrict Gal4-dependent Kr RNAi induction to specific developmental windows. At 19°C, Gal80ts suppresses Gal4 activity and inhibits transgene expression, whereas shifting animals to 29°C inactivates Gal80ts and induces Kr RNAi expression. Animals were shifted between 19°C and 29°C at the embryonic, larval, pupal and adult stages according to the experimental scheme shown in Fig. 3C. Adult brains from each condition were dissected after EdU feeding, fixed and stained for EdU, Mira and DAPI as described above. EdU-positive MBNB clones were manually counted per brain hemisphere.

### Quantification of Kr signal intensity in MBNBs during pupal development (Fig. 3F, S3F)

Pupae were collected over a 24 h period and subsequently incubated for 0–24, 24–48, 48–72 or 72–96 h after puparium formation (APF) before brain dissection. Pupal brains were fixed and immunostained as described above using rat anti-Kr and rabbit anti-Kr antibodies. Z-stack images encompassing the MB cell body region were acquired from the posterior side using a Nikon C2 confocal microscope. MBNBs were identified based on their stereotypical dorsoposterior location, large rounded morphology, low DAPI signal, proximity to the developing mushroom body calyx and, where applicable, the surrounding lineage structure.

Kr signal intensity was quantified in individual MBNBs using ImageJ/Fiji. DAPI images were used to define nuclear and cytoplasmic regions. For each MBNB, mean Kr signal intensity was measured separately in the nuclear and cytoplasmic regions, and local background was measured from neighbouring brain tissue outside the MBNB. Normalised Kr signal intensity was calculated as (mean ROI intensity − background intensity) / background intensity and plotted for each pupal stage.

To validate the specificity of the Kr antibody signal in MBNBs, 24–48 h APF pupal brains from control and *insc>KrIR* animals expressing *insc*>GFP were fixed and immunostained with rat anti-Kr antibody. Z-stacks covering the MB cell body region and part of the MB calyx were acquired at 1.5 µm intervals in three channels: DAPI, GFP and Alexa Fluor 568 for Kr. MBNBs were identified using *i<u>nsc</u>*>GFP signal, DAPI morphology and position within the MB cell body region. Cytoplasmic Kr signal intensity was measured in individual MBNBs, with local background measured from adjacent brain tissue outside the MBNB. Normalised Kr signal intensity was calculated as (mean MBNB cytoplasmic intensity − background intensity) / background intensity.

### Quantification of Imp- and Syp-positive areas within MB cell body regions in adult brains (Fig. 4A)

Adult brains were dissected, fixed and immunostained as described above. Brains were stained with rat anti-Imp, rabbit anti-Syp and mouse anti-Eyeless (Ey) antibodies, followed by appropriate Alexa Fluor-conjugated secondary antibodies. Ey signal was used to define the MB cell body region. Brains were mounted with the posterior side facing the coverslip and imaged using a Nikon C2 confocal microscope. Z-stacks covering the MB cell body region and part of the MB calyx were acquired at 1 µm intervals in four channels: DAPI, Alexa Fluor 488 for Syp, Alexa Fluor 568 for Imp and Alexa Fluor 647 for Ey.

For quantification, 3D reconstructions were generated using the Volume View/Max Intensity Projection function in Nikon NIS-Elements. Reconstructed images were rotated to present the dorsoposterior MB cell body surface region facing the viewer, and the same orientation strategy was applied to all samples. Imp and Syp signals were thresholded using equivalent settings within each experiment to generate binary masks for area measurement. The Ey-positive MB cell body surface region was defined as the reference area, and the Imp- or Syp-positive area within this region was measured using the Area function in NIS-Elements. The ratio of Imp- or Syp-positive area to the total Ey-positive MB cell body surface area was calculated for each hemisphere.

### Quantification of Imp and Syp signal intensity in MBNBs in pupal brains (Fig. 4B)

Pupal brains at 72–96 h APF were dissected, fixed and immunostained as described above. MBNBs were visualised using *insc-*Gal4-driven UAS-mCD8::GFP. Brains were stained with rat anti-Imp and rabbit anti-Syp antibodies, followed by appropriate Alexa Fluor-conjugated secondary antibodies. Native GFP fluorescence from UAS-mCD8::GFP was used to identify MBNBs. Brains were mounted with the posterior side facing the coverslip and imaged using a Nikon C2 confocal microscope. Z-stacks covering the MB cell body region and part of the MB calyx were acquired at 1.5 µm intervals in four channels: DAPI, GFP, Alexa Fluor 568 for Imp and Alexa Fluor 647 for Syp. MBNB regions of interest were defined based on *insc>*GFP signal, cell size and position within the MB cell body region. For each MBNB, mean Imp or Syp fluorescence intensity was measured, and local background intensity was measured from adjacent brain tissue outside the MBNB. Normalised signal intensity was calculated as (mean MBNB fluorescence intensity − background intensity) / background intensity.

### Developmental expression profiles of temporal factors in the MBNB lineage (Fig. S5A)

Developmental expression profiles of temporal regulators in the MBNB lineage were analysed using the MB lineage-specific RNA-seq dataset reported by Liu et al. (27) Mean expression values ± SD from three biological replicates were extracted for selected temporal factors at 24 h ALH, 50 h ALH, 84 h ALH and 36 h APF, and are presented in Fig. S5A.

### Quantification of E93 and Kr-h1::GFP signals (Fig. 5B, S5B)

Pupal brains at 72–96 h APF were dissected, fixed and immunostained as described above. Brains were stained with guinea pig anti-E93, chicken anti-GFP and rat anti-Mira antibodies, followed by appropriate Alexa Fluor-conjugated secondary antibodies. Kr-h1::GFP was visualised using anti-GFP staining, E93 using Alexa Fluor 568, and Mira using Alexa Fluor 647. Brains were mounted with the posterior side facing the coverslip and imaged using a Nikon C2 confocal microscope. Z-stacks covering the MB cell body region and part of the MB calyx were acquired at 1.5 µm intervals.

For each MBNB, a maximum-intensity projection was generated from the confocal z-sections containing the MBNB and its associated lineage. MBNBs were identified based on Mira staining, cell morphology and position within the MB cell body region. An MBNB ROI was defined around the Mira-positive MBNB. A second ROI encompassing the surrounding MBNB lineage/progeny region was defined around the neighbouring progeny region, excluding the MBNB ROI. For Fig. 5B, E93 fluorescence intensity was quantified separately in the MBNB ROI and the surrounding progeny ROI. Kr-h1::GFP fluorescence intensity was quantified only in the MBNB ROI. For Fig. S5B, E93 signal intensity was quantified within the insc>GFP-positive MB lineage region.

Local background fluorescence was measured from nearby brain tissue outside the mushroom body lineage region, avoiding obvious E93-positive cells and regions outside the brain tissue. Normalised fluorescence intensity was calculated as (ROI mean intensity − background mean intensity) / background mean intensity. Raw ROI mean intensity, background intensity and background-subtracted intensity values were retained for quality control.

### Differential expression analysis of cell cycle genes in the *Drosophila* adult and larval brains (Fig. 1B, C, S1)

The original data files of the single-cell RNA sequencing of *Drosophila* adult and larval brains (47) were obtained from the NCBI Gene Expression Omnibus (series number GSE107451) and processed using Seurat v3 (79). To ensure data quality and remove potential dying cells, we restricted the dataset to samples with 200 to 3700 unique genes and a UMI-to-gene ratio smaller than 6. The resulting dataset comprised 56,619 samples and 12,925 features. For normalisation, raw counts were first adjusted to account for sequencing depth by scaling each cell by the total number of counts and multiplying by a scaling factor of 10,000. The normalised counts were then log-transformed using the natural logarithm (log1p), which transforms the data using the formula log(1 + x), where x represents the normalised counts. Principal Component Analysis (PCA) was conducted on the 2000 highly variable genes identified. Shared Nearest Neighbour (SNN) cell clustering at a resolution of 0.2 or 8 was performed using the first 20 principal components, and t-SNE reduction was applied to visualise the clustering results. Based on the differential expression of the cluster markers and comparison with the original annotations (47), 17 or 84 cell clusters were identified, respectively.

The list of 112 CCR genes was manually curated from *Drosophila* orthologues and known regulators of cell-cycle control, including core cell-cycle regulators, cyclins and CDKs, DNA replication factors, checkpoint regulators, mitotic regulators, APC/C and ubiquitin-proteasome components, and non-canonical CDKs with known transcriptional roles. The expression of these CCR genes across the identified clusters was then visualised using dot plots. In these plots, the colour intensity of each dot represents the average expression level of the CCR gene, and the size of each dot indicates the proportion of cells within the cluster expressing the gene.

For the analysis of *Drosophila* larval brains, the original data files of the single-cell RNA sequencing of larval brains (48) were obtained, processed and presented similar to the adult brain. The analysis dataset contains 4349 samples and 12,942 features. A resolution of 2 was used and 29 clusters were identified.

### Cell cycle regulator overexpression in postmitotic neurons and survival analysis (Fig. S2A-C)

To test whether forced expression of positive cell cycle regulators can drive postmitotic neurons into the cell cycle, E2F1-Dp and Cdk2-CycE were overexpressed in dopaminergic neurons using TH-Gal4 or in mushroom body neurons using mb247-Gal4. For adult induction experiments, young adult flies were shifted to 29°C to induce Gal4-dependent transgene expression and maintained for 10 days before brain dissection. For late pupal induction experiments, animals carrying Gal80ts were shifted to 29°C approximately two days before eclosion. Brains were dissected, fixed and stained for pH3 to detect mitotic entry, Dcp-1 to detect apoptosis where indicated, GFP reporter expression and DAPI. pH3-positive cells were manually scored in the relevant neuronal populations. For experiments testing age-dependent refractoriness to cell cycle re-entry, gene induction was initiated in older adult flies 10 days after eclosion.

For survival analysis, E2F1-Dp and Cdk2-CycE were overexpressed in dopaminergic neurons using *TH*-Gal4 combined with Gal80ts. Experimental flies carried *TH-Gal4, UAS-Dp-E2F1, UAS-CycE-Cdk2, UAS-mCD8::GFP* and *tub-Gal80ts*, whereas control flies expressed *lacZ* under the same driver system. Gene expression was induced by shifting 2-day-old adult flies from 19°C to 29°C. Survival was monitored daily, and Kaplan–Meier survival curves were generated from three independent biological replicates.

### Statistical analysis

Statistical analyses were performed using GraphPad Prism v10.1.2. The statistical tests used for each experiment are indicated in the corresponding figure legends. Unless otherwise stated, tests were two-sided. Sample sizes are indicated in the figures or figure legends and refer to the number of brain hemispheres, MBNBs, cells or flies analysed, as appropriate. Data are presented as mean ± SD, except for bounded count data analysed using non-parametric tests, which are presented as median and interquartile range where indicated. Statistical significance is indicated as follows: *p ≤ 0.05, **p ≤ 0.01, ***p ≤ 0.001, ****p < 0.0001; ns, not significant.

## Acknowledgements

We thank Dr Margaret Ho for discussion, for sharing *Drosophila* reagents and for providing the initial training for the dissection and immunostaining of *Drosophila* adult brains, Dr Wei Wu at Shanghai Fly Center, Dr Simon Collier at Cambridge *Drosophila* Facility at ShanghaiTech University for the help with maintenance and transporting *Drosophila* stocks and for embryo injection services, Dr. Reinhard Klug and the VDRC for the assistance with the transportation of *Drosophila* stocks and RNAi fly lines, and Drs Cédric Maurange, Cahir O’Kane, Kei Ito, Nan Liu, Yukinori Hirano, and Torcato Martins, for discussion, Ilan Davis, Chris Rushlow, Claude Desplan, Kei Ito, Kuniaki Saitoh, Rita Sousa-Nunes, Hua Bai, Kang Peng, and Chenyu He for antibodies and technical advice, and all *Drosophila* colleagues at ShanghaiTech University, in particular, Drs Ji-long Liu, Jingnan Liu, Guanjun Guo, and Chenhui Wang, and China *Drosophila* community for discussion, sharing information and reagents. We also thank all Kimata lab members at ShanghaiTech University and the University of Cambridge for cooperation and discussion. This work was supported by the ShanghaiTech University start-up grant (2018F0202-000-06), the National Natural Science Foundation of China (NSFC) Mianshang Project (32170746), and the NSFC Research Fund for International Scientists (RFIS, 32150710520) awarded to Y.K.

## Supplementary figure legends

**Figure S1.**
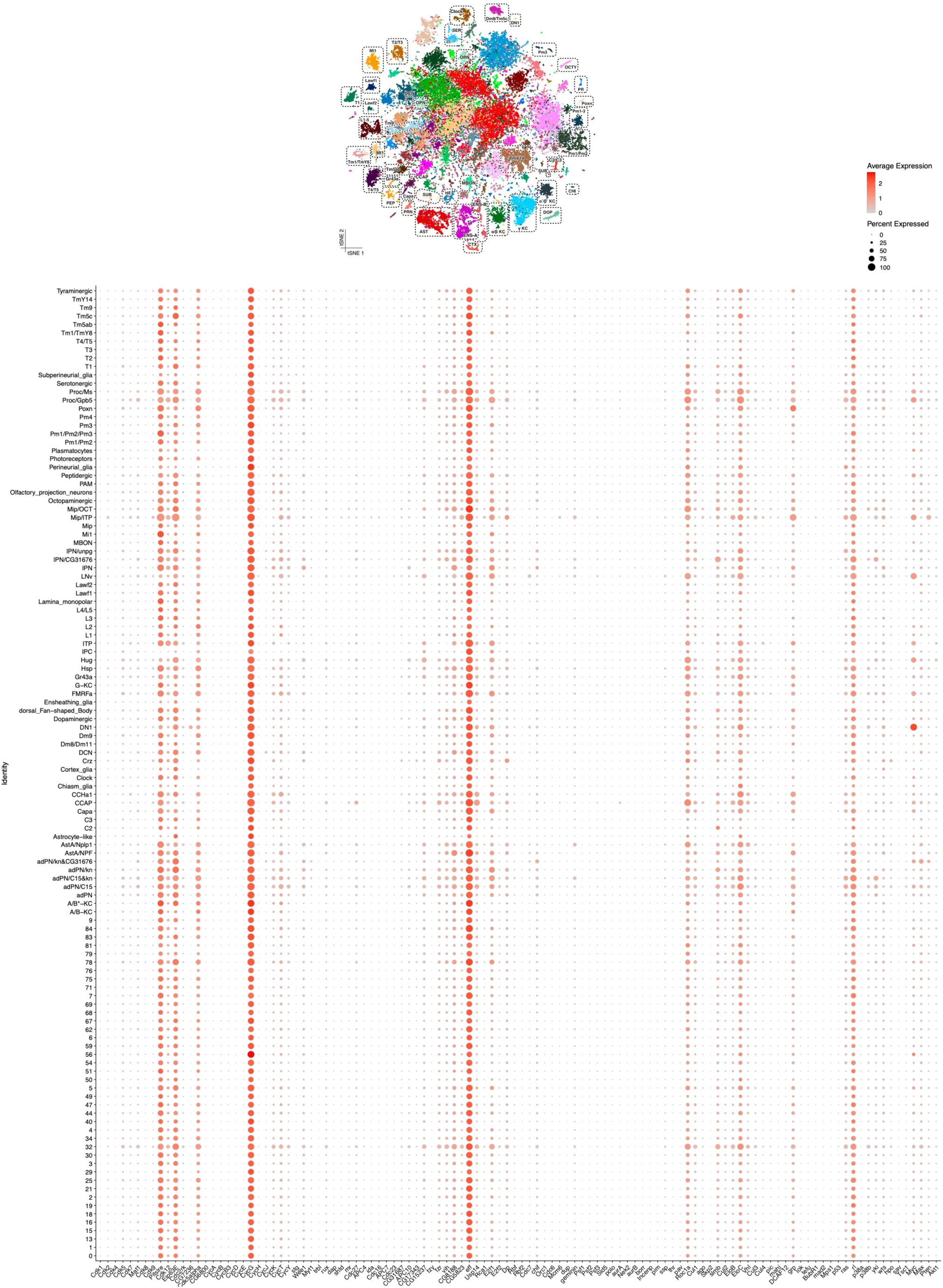
Expression of CCR genes across 84 cell clusters in the adult *Drosophila* brain. Top: tSNE plot showing 84 cell clusters identified in the adult *Drosophila* brain scRNA-seq data (Davie et al., 2018). Bottom: dot plot showing expression levels of 112 selected CCR genes across the 84 clusters. Colour intensity indicates average expression level, and dot size indicates the percentage of cells within each cluster expressing the gene. Positive CCRs, including *Cdk1*, *Cdk2*, *CycB* and *Polo*, are largely absent from neuronal and glial clusters but are enriched in a small neuroblast-like population. In contrast, negative regulators such as *fzr*, *Wee1* and *rux* remain broadly expressed across neuronal clusters, consistent with maintenance of the postmitotic state in the adult brain.

**Figure S2.**
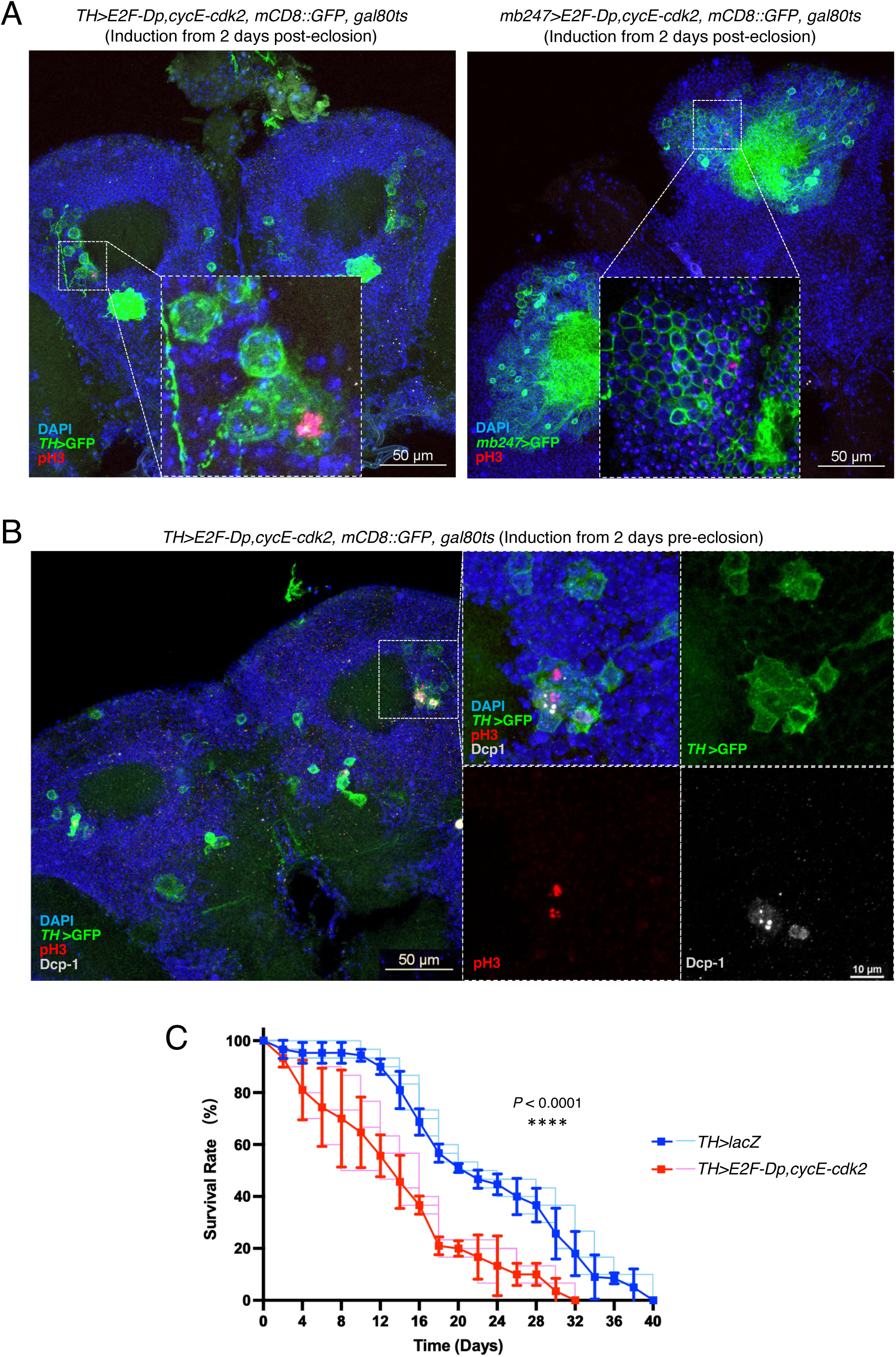
Forced expression of positive CCRs induces mitotic entry in postmitotic neurons. **(A)** Overexpression of Dp-E2F1 and CycE-Cdk2 in dopaminergic neurons using *TH-*Gal4 (left panels) and in mushroom body neurons using *mb247-*Gal4 (right panels) induces mitotic entry in young adult flies, ∼2 days post-eclosion. Gene expression was induced by shifting flies to 29°C for 10 days. Brains were stained for DAPI (blue), GFP reporter expression (green) and pH3 (red). pH3-positive neurons were detected following E2F1-Dp and Cdk2-CycE overexpression, whereas no pH3-positive neurons were observed in controls. When gene induction was initiated in older flies, 10 days post-eclosion, no pH3-positive neurons were detected (*n* ≥ 10), suggesting that neurons become increasingly refractory to cell-cycle re-entry with age. Scale bars: 50 μm. **(B)** Overexpression of Dp-E2F1 and CycE-Cdk2 in dopaminergic neurons during late pupal development (∼2 days before eclosion) allowed some adult flies to emerge; however, these flies exhibited premature lethality, dying within 3–5 days after eclosion. Brains dissected 5 days after gene induction contained pH3-positive neurons (12 out of 27 brains examined). Some of these neurons also showed Dcp-1 staining (grey), indicating apoptosis. Overexpression of the same CCR combination in the mushroom body neurons using *mb247-*Gal4 caused lethality before adult eclosion. Scale bars: 50 μm. **(C)** Kaplan-Meier survival analysis of adult flies overexpressing Dp-E2F1 and CycE-Cdk2 in dopaminergic neurons. Experimental flies (*TH>E2F-Dp, CycE-Cdk2, mCD8::GFP, gal80ts*) and controls (*TH>lacZ, mCD8::GFP, gal80ts*) were shifted from 19°C to 29°C two days after eclosion to induce transgene expression. Flies overexpressing E2F1-Dp and Cdk2-CycE exhibited significantly reduced lifespan compared to controls (*P* < 0.0001, log-rank Mantel-Cox test). Pale blue and pale red lines represent three individual biological replicates, and dark blue and red lines represent mean survival curves. Error bars indicate SD.

**Figure S3.**
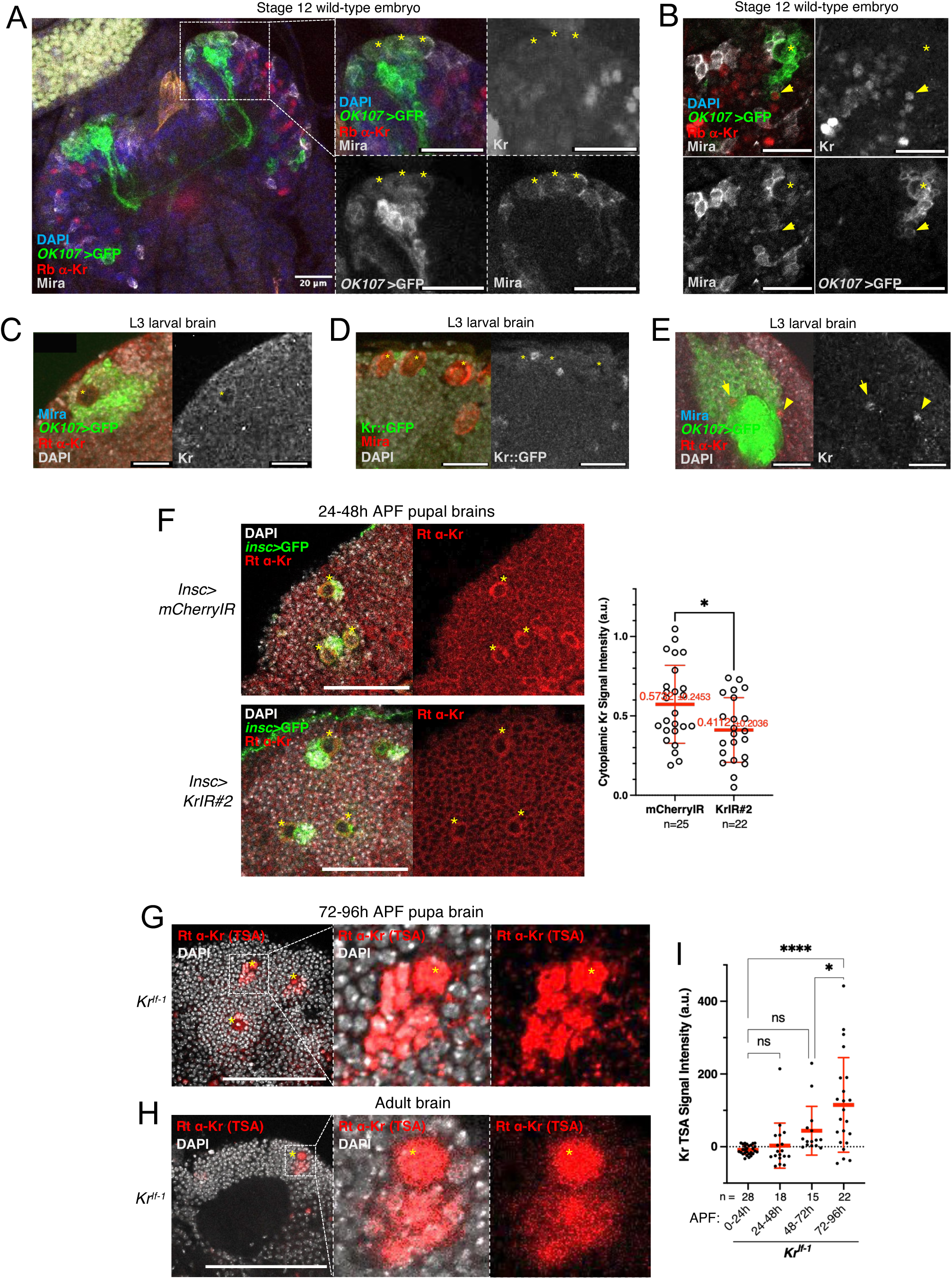
Kr expression in MBNBs during development. **(A, B)** Representative confocal images of stage 12 wild-type embryonic central nervous system (CNS) stained with rabbit anti-Kr antibody (red), *OK107*>GFP (green), DAPI (blue), and Mira (grey). (A) Overview of the embryonic CNS and magnified views of the MB lineage region, with merged images and individual greyscale channels shown. Yellow asterisks indicate MBNBs identified by *OK107*>GFP and Mira co-expression. (B) Representative images of the MB lineage region. Yellow asterisks indicate MBNBs, and yellow arrows indicate a neuron within the MB lineage expressing Kr. Although Kr was broadly expressed in the embryonic CNS, it was not detected in MBNBs. Scale bars: 20 µm. **(C-E)** Kr expression in third-instar larval brains. (C) Confocal image of the MB cell body region stained for *OK107*>GFP (green), Mira (cyan), and Kr (red) using a rat anti-Kr antibody. The individual Kr channel is shown in grey. Yellow asterisks mark MBNBs, which exhibit weak cytoplasmic Kr signals. (D) Kr::GFP(Bac) reporter expression in the MB cell body region, showing Kr::GFP (green), DAPI (grey), and Mira (red). The individual GFP channel is shown in grey. Asterisks mark MBNBs. Strong Kr::GFP signals were detected in single cells adjacent to MBNBs, likely corresponding to GMCs or early-born neurons that may have inherited Kr::GFP from MBNBs. (E) Confocal image of the MB calyx region stained for *OK107*>GFP (green), Mira (cyan) and Kr (red), with the individual Kr channel shown in grey. Kr-expressing neurons near the MB calyx are indicated by arrowheads. Scale bars: 20 µm. **(F)** Kr antibody signal is reduced following Kr depletion. Left: representative confocal images of 24-48 h APF pupal brains from control and *insc>KrIR#2* conditions, stained with rat anti-Kr antibody (red), DAPI (grey) and *insc>*GFP (green). Scale bar: 50μm. Right: quantification of normalised cytoplasmic Kr signal intensity. Scatter dot plot shows individual MBNB measurements. Thick and thin red bars indicate means and SDs, respectively. Statistical significance was determined using an unpaired t-test. See Materials and methods for details on quantification. **(G, H)** Representative confocal images of *Kr^If-1^* mutant late pupal (G) and adult (H) brains stained with rat anti-Kr antibody (red) and DAPI (grey) using tyramide signal amplification (TSA). Yellow asterisks indicate MBNBs identified by their position and DAPI morphology. Scale bars: 100 µm. **(I)** Quantification of TSA-enhanced Kr signal intensities in individual MBNBs in *Kr^If-1^*mutant pupal brains, showing Kr misexpression in MBNBs during late pupal stages in *Kr^If-1^* mutants. Statistical significance is indicated as follows: *p ≤ 0.05, **p ≤ 0.01, ***p ≤ 0.001, ****p < 0.0001; ns, not significant.

**Figure S4.**
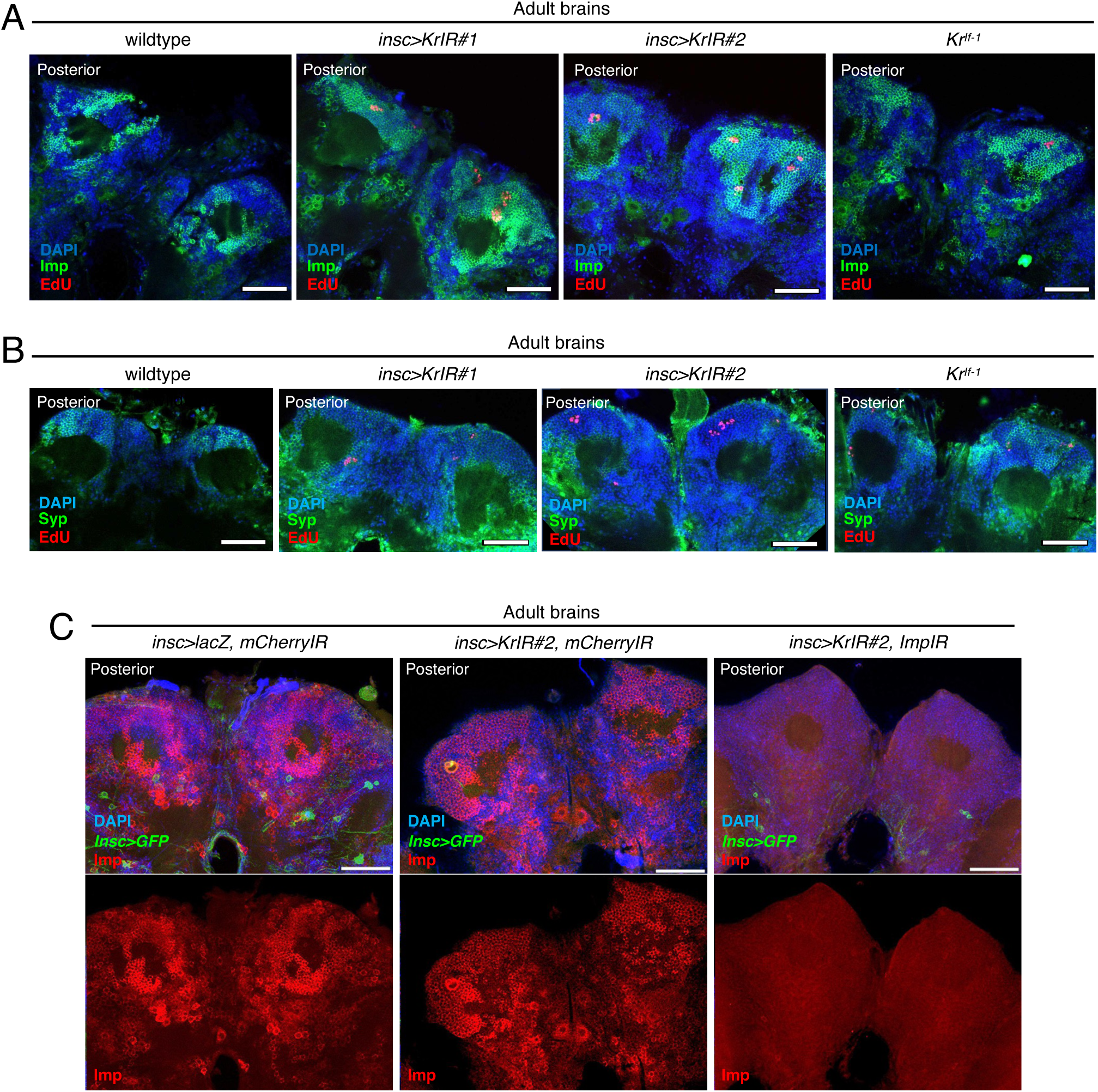
Imp expression persists in MBNBs in *Kr* RNAi and *Kr^If-1^* mutant flies. **(A)** Representative confocal images of dorsoposterior regions of wild-type, *Kr^If-1^* mutant and *insc>KrIR* adult brains. EdU labelling (red) marks proliferating cells, Imp expression (green) was visualised using an Imp-specific antibody, and DAPI (blue) marks DNA. Compared to controls, Imp-expressing regions around the MB cell body region were expanded in *Kr^If-1^* and *insc>KrIR* brains. Scale bars: 50 µm. **(B)** Representative confocal images of dorsoposterior MB cell body regions in wild-type, *Kr^If-1^* and *insc>KrIR* adult brains. EdU labelling (red) marks proliferating cells, Syp expression (green) was visualised using Syp-specific antibodies, and DAPI (blue) marks DNA. These images correspond to the Syp analysis shown in Fig. 4A. Scale bars: 50 µm. **(C)** Validation of Imp depletion in Kr RNAi adult brains. Representative confocal images of control *insc>mCherryIR*, *insc>KrIR#2, mCherryIR* and *insc>KrIR#2, ImpIR* adult brains stained for Imp (red), i*nsc*>GFP (green) and DAPI (blue). Expression of *Imp* RNAi under *insc-*Gal4 efficiently depleted endogenous Imp proteins in brain tissue, including the MB lineage region. Scale bars: 50 µm.

**Figure S5.**
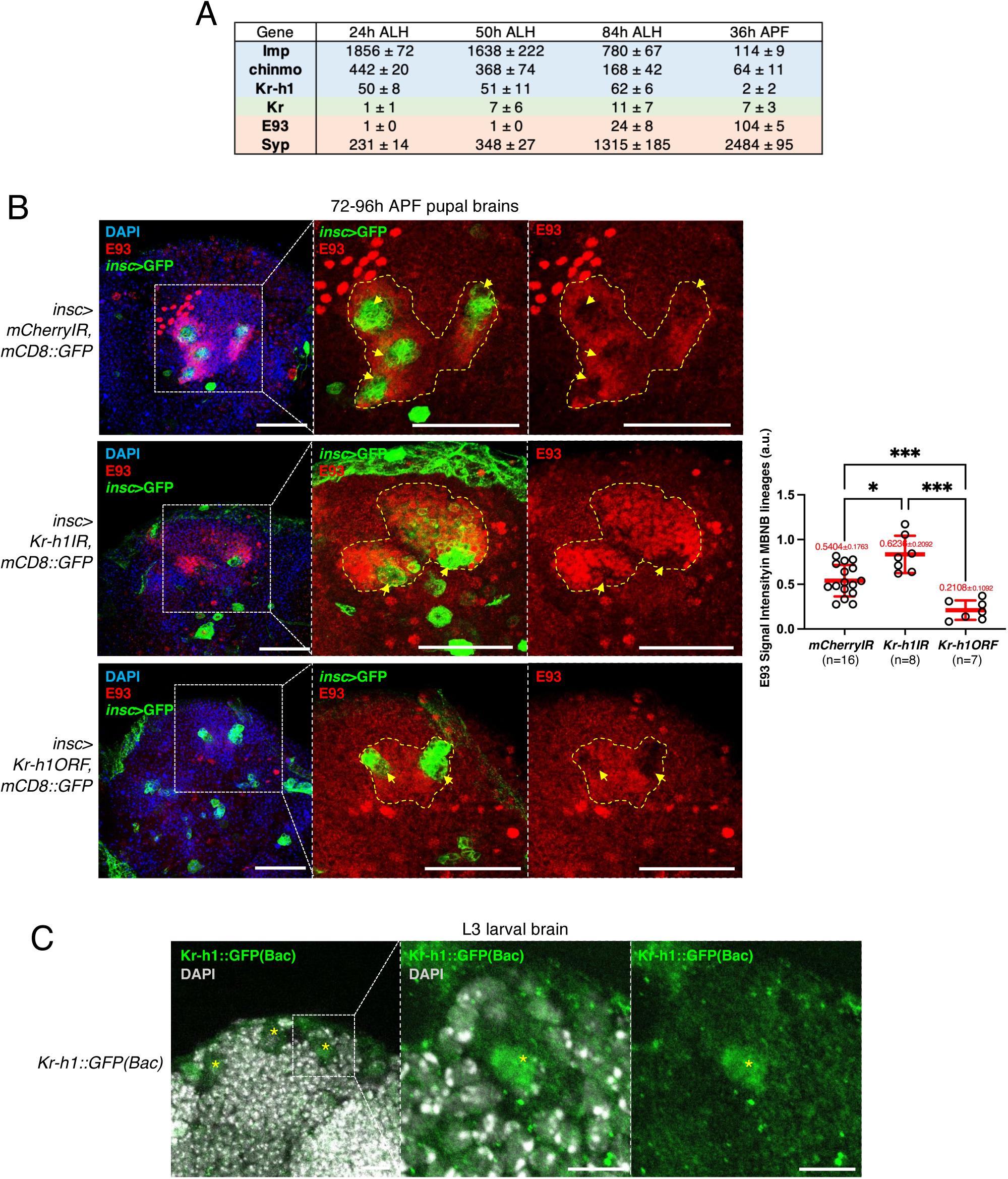
E93 expression in the MB lineages is regulated by Kr-h1. **(A)** Developmental expression profiles of temporal factors in the MBNB lineage. Expression levels (mean ± SD) of selected temporal regulators in MB lineages at 24 h ALH, 50 h ALH, 84 h ALH and 36 h APF are shown. Values were calculated from three biological replicates using the MB lineage-specific RNA-seq dataset reported by Liu et al. (2015). Imp, chinmo and Kr-h1 are highly expressed during larval stages and decline during pupal development, whereas Syp and E93 are strongly induced during the larval-to-pupal transition. Kr expression remains relatively low throughout development. **(B)** Kr-h1 regulates E93 expression in the MB lineage during late pupal development. Left: representative confocal images of 72–96 h APF pupal brains from control *insc>mCherryIR, mCherryIR*, *insc>Kr-h1IR, mCherryIR* and *insc>Kr-h1ORF::FLAG, mCherryIR* flies, stained for E93 (red) and *insc*>GFP (green). Dotted lines indicate *insc>*GFP-positive MB lineage regions. Scale bars: 50 µm. Right: scatter dot plots showing normalised E93 signal intensity in the MB lineage region, quantified per hemisphere. Thick and thin red bars indicate means and SDs, respectively. n indicates the number of hemispheres analysed per condition. Statistical significance was determined using pairwise Mann–Whitney U tests. *p ≤ 0.05, **p ≤ 0.01, ***p ≤ 0.001, ****p < 0.0001; ns, not significant. **(C)** Representative confocal images showing Kr-h1::GFP(Bac) expression in MBNBs in third-instar larval brains. Kr-h1::GFP was enriched in the nuclei of MBNBs. Scale bars: 10 µm.

## Notes

### Competing Interest Statement

The authors have declared no competing interest.

### Summary of Updates

This revised version has been updated to correspond to the revised manuscript submitted to eLife. The revision includes new analyses and experimental data addressing Kr expression in the KrIf-1 background, Imp/Syp regulation, Kr-h1/E93 regulation, and an updated model for MBNB termination.

